# FUS-dependent liquid-liquid phase separation is an early event in double-strand break repair

**DOI:** 10.1101/798884

**Authors:** Brunno R. Levone, Silvia C. Lenzken, Marco Antonaci, Andreas Maiser, Alexander Rapp, Francesca Conte, Stefan Reber, Antonella E. Ronchi, Oliver Mühlemann, Heinrich Leonhardt, M. Cristina Cardoso, Marc-David Ruepp, Silvia M.L. Barabino

**Author notes:** Corresponding author: Silvia M.L. Barabino, Department of Biotechnology and Biosciences, University of Milano-Bicocca, Piazza della Scienza, 2, I-20126 Milano, Italy. These two authors contributed equally to this work (Andreas Maiser, Alexander Rapp). Institute of Molecular Biology (IMB), Ackermannweg 4, 55128 Mainz, Germany.

## Abstract

RNA-binding proteins (RBPs) are emerging as important effectors of the cellular DNA damage response (DDR). The RBP FUS is implicated in RNA metabolism and DNA repair, and it undergoes reversible liquid-liquid phase separation (LLPS) *in vitro*. Here, we demonstrate that FUS-dependent LLPS is necessary for the initiation of the DDR. Using laser microirradiation in FUS-knockout cells, we show that FUS is required for the recruitment to DNA damage sites of the DDR factors KU80, NBS1, 53BP1, and of SFPQ, another RBP implicated in the DDR. The relocation of KU80, NBS1, and SFPQ is similarly impaired by LLPS inhibitors, or LLPS-deficient FUS variants. We also show that LLPS is necessary for efficient γH2AX foci formation. Finally, using super-resolution structured illumination microscopy, we demonstrate that the absence of FUS impairs the proper arrangement of γH2AX nano-foci into higher-order clusters. These findings demonstrate the early requirement for FUS-dependent LLPS in the activation of the DDR and the proper assembly of DSBs repair complexes.

## Introduction

DNA damage is a threat for cell survival. Unrepaired DNA damage can lead to genome instability, a hallmark of cancer cells. To counteract DNA damage, cells have evolved a complex cellular response, commonly referred to as DNA damage response (DDR). The early DDR events have been best elucidated at sites of DNA double strand breaks (DSBs), which are the most dangerous type of DNA lesions. After the occurrence of a DSB, the sensor protein kinases DNA-PK (DNA-dependent protein kinase), ATM (ataxia telangiectasia mutated), and ATR (ATM and Rad3 related) are rapidly activated, and the KU70/KU80 (XRCC6/XRCC5) heterodimer is recruited to the broken DNA ends. Phosphorylation of the histone variant H2AX at S139 (known as γH2AX) by ATM serves as an early mark of DNA damage and as a platform for the recruitment of early DDR factors such as MDC1, and the MRN complex (consisting of MRE11, NBS1 and RAD50) (Stucki and Jackson, 2006). In mammalian cells, the accumulation of DDR factors at sites of DNA damage gives rise to subnuclear foci that can be readily visualised microscopically. Using super resolution microscopy, others and we have recently shown that these foci correspond to clusters of nano-domains, and this clustering is required for an efficient DNA damage repair (Lopez Perez et al., 2016; Natale et al., 2017). In mammals, DSBs are eventually repaired via two main pathways, depending on the cell cycle phase: homologous recombination (HR), which repairs DSBs in late S and G2 phases, and non-homologous end joining (NHEJ), which is active throughout the entire cell cycle.

In addition to canonical DDR factors, large-scale proteomic and genomic studies have identified several RNA-binding proteins (RBPs) as potential novel DDR factors, either as targets of the apical DDR kinases (Matsuoka et al., 2007) or as proteins that, when absent, lead to the activation of the DDR (Paulsen et al., 2009). RBPs have been shown to contribute both directly and indirectly to genome stability. For example, the loss of pre-mRNA splicing or mRNA export factors can favour the accumulation of RNA:DNA hybrids (R-loops) that can be processed to DSBs (Chuang et al., 2019; Li and Manley, 2005). In addition, an increasing number of studies have shown that several RBPs are recruited to sites of DNA damage and participate in DSB repair (Mikolaskova et al., 2018).

The multifunctional DNA/RNA-binding protein Fused in Sarcoma (FUS) is involved in splicing, translation, and mRNA transport (Dormann and Haass, 2013). FUS is a member of the FET family of proteins that also includes Ewing’s sarcoma (EWS) and TATA-binding protein-associated factor 2N (TAF15). Both *in vivo* and *in vitro* observations point towards a role for FUS in maintaining genome stability. Mice lacking FUS are hypersensitive to ionising radiation (IR), show defects in spermatogenesis, and chromosomal instability (Hicks et al., 2000; Kuroda et al., 2000). *In vitro*, FUS stimulates the formation of DNA loops (D-loops) between complementary DNA molecules, structures that correspond to one of the first steps in HR (Baechtold et al., 1999). In cells, FUS is recruited very early to sites of DNA damage (Aleksandrov et al., 2018; Mastrocola et al., 2013) and its silencing leads to an impairment of DSB repair both by HR and NHEJ (Mastrocola et al., 2013; Wang et al., 2013). In addition, FUS is an ATM and DNA-PK substrate (Deng et al., 2014; Gardiner et al., 2008).

The N-terminal region of FUS (residues 1–165) is a highly conserved prion-like domain (PLD) composed primarily of serine, tyrosine, glycine, and glutamine (QGSY-rich). This domain mediates protein-protein interactions and drives the aggregation of FUS into protein inclusions (Sun et al., 2011). Several studies have shown that the PLD of FUS undergoes a reversible dynamic phase transition between a disperse state, liquid droplets, and hydrogels (Kato et al., 2012; Murakami et al., 2015; Patel et al., 2015). FUS liquid-liquid phase separation (LLPS) occurs both *in vivo* and *in vitro* at physiological concentrations (Burke et al., 2015; Murakami et al., 2015).

It is increasingly recognised that LLPS provides a molecular basis for the formation of subcellular membrane-less organelles such as nucleoli, Cajal bodies, paraspeckles, and stress granules (Boeynaems et al., 2018). Paraspeckles are subnuclear compartments of poorly characterised function that assemble on the lncRNA NEAT1, which induces phase separation of four core RBPs: SFPQ, NONO, FUS and RBM14 (Hirose et al., 2019). Interestingly, all these proteins contain PLDs of variable length (Harrison and Shorter, 2017). The PLDs of FUS and RBM14 are required for *in vitro* phase separation and *in vivo* paraspeckle formation (Hennig et al., 2015). SFPQ, its paralog NONO, and RBM14 are also RBPs implicated in DNA repair (Kai, 2016). SFPQ silencing was reported to sensitise cells to DNA crosslinking and alkylating agents, and to reduce DSB repair by HR (Rajesh et al. 2011). SFPQ has DNA re-annealing and strand-invasion activity that may lead to the formation of D-loop structures (Akhmedov and Lopez, 2000). In addition, SFPQ, NONO, and RBM14 also promote NHEJ (Bladen et al., 2005; Jaafar et al., 2017; Simon et al., 2017).

Despite all the observations implicating RBPs such as FUS and SFPQ in DNA damage repair, their precise molecular function remains to be fully elucidated.

Here, we show that FUS is required for the recruitment of SFPQ and the retention of KU80 on DSBs. Consistent with recent results indicating a role for LLPS in DNA damage repair (Kilic et al., 2019; Pessina et al., 2019), we show that LLPS inhibitors impair the formation of γH2AX and 53BP1 foci and the proper recruitment of FUS, SFPQ, KU80 and 53BP1. Moreover, LLPS-deficient variants of FUS affect accumulation of SFPQ at sites of laser-induced DNA damage. Finally, we demonstrate that FUS is needed for the higher order clustering of γH2AX chromatin nano-domains, which is required for efficient DNA damage repair (Natale et al., 2017). Overall, our findings highlight an important physical rather than enzymatic role of FUS in promoting LLPS at DNA damage sites and the recruitment of early key DDR factors.

## Results

### FUS knockout sensitises human cells to DNA damage

To shed light on the role of FUS in DDR and genome stability, we generated human HeLa and SH-SY5Y cells in which the FUS gene was knocked-out (FUS-KO) by CRISPR-trap genome editing technique (Reber et al., 2018). Western blot analysis showed a substantial increase in the level of endogenous γH2AX both in HeLa FUS-KO and in SH-SY5Y FUS-KO cells (Figure 1A and 1B, and Suppl. Figure 1A and 1B, respectively). Similarly, also the shRNA-mediated knockdown of FUS in SH-SY5Y cells was sufficient to increase H2AX phosphorylation (data not shown). Consistently, immunofluorescence analyses revealed increased formation of γH2AX foci both in HeLa FUS-KO and in SH-SY5Y FUS-KO cells (Figure 1C and 1D, and Suppl. Figure 1D and 1E, respectively). These observations suggest that, in the absence of FUS, cells either generate more DNA damage or repair DNA damage less efficiently. Indeed, transient transfection of HeLa FUS-KO cells with exogenous Flag-tagged FUS reduced the number of γH2AX foci (Figure 1E).

**Figure 1.**
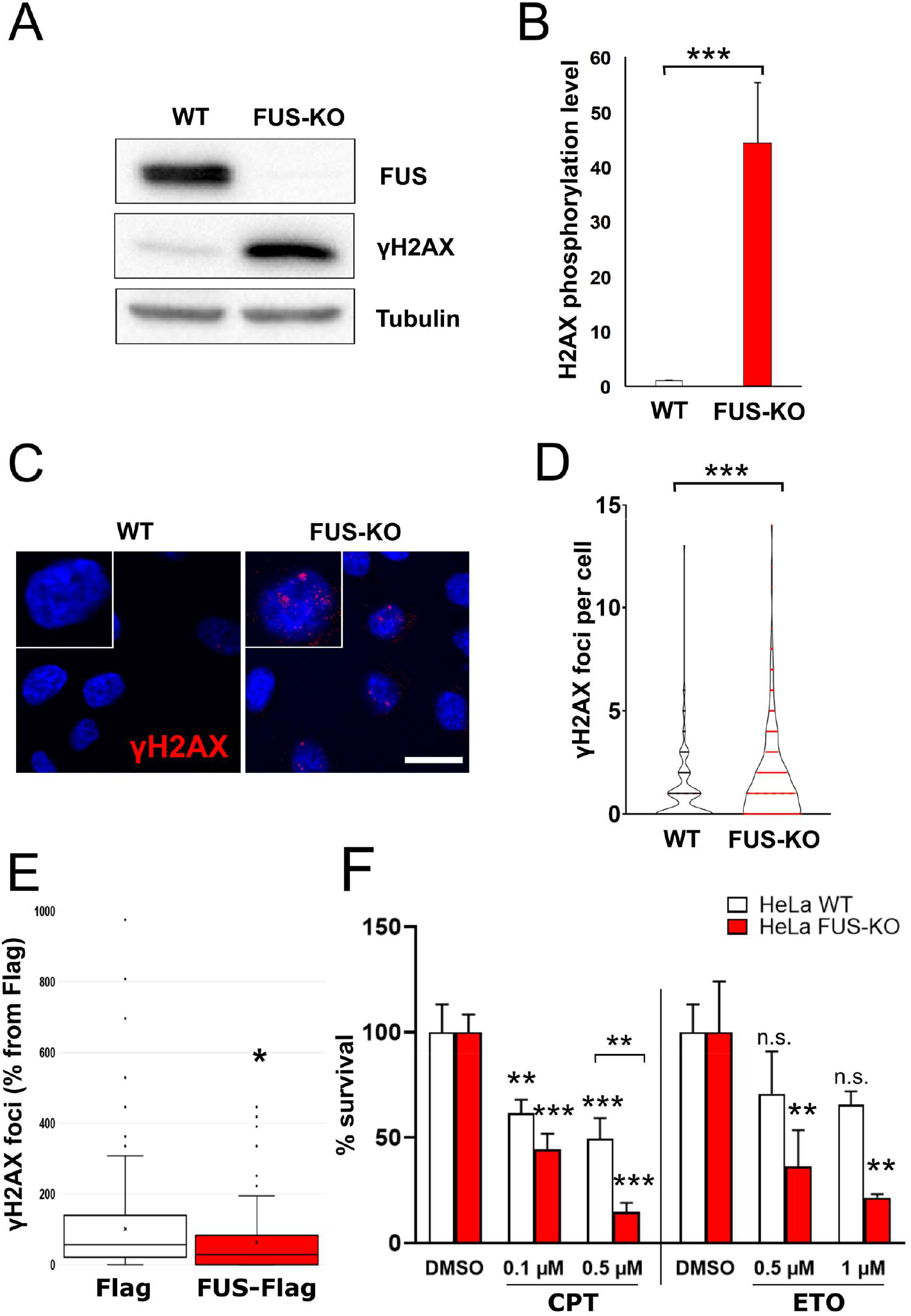
Loss of FUS results in accumulation of DNA damage and sensitisation to genotoxic insult in HeLa cells. A. Total extracts of wild type and FUS-KO HeLa cells were analysed by Western blotting with anti-FUS and anti-γH2AX antibodies. Tubulin was used as loading control. B. Quantification of γH2AX protein level. Statistics: Student’s t-test. C. Representative confocal micrographs of γH2AX foci in wild type and FUS-KO HeLa cells. Cropped single cells are 2x enlarged. Scale bar: 20 µm. D. Quantification of γH2AX foci. The number of foci per nucleus was counted using ImageJ. Data from two biological replicates, with 170 cells per replicate. Statistics: Student’s t-test. E. HeLa FUS-KO cells were transiently transfected with a plasmid expressing Flag-tagged FUS and stained with anti-Flag and anti-γH2AX. Foci were quantified by ImageJ. Statistics: Student’s t-test. F. HeLa wild type and FUS-KO cell viability assessed by Trypan blue staining upon treatment with increasing concentrations of camptothecin (CPT, 0.1 or 0.5 µM) or etoposide (ETO, 0.5 or 1 µM). Statistics: two-way ANOVA followed by Bonferroni post-hoc test In all panels: n.s., non-significant; *, p<0.05, **, p < 0.01 and ***, p < 0.001.

Next, we investigated whether the knockout of FUS results in DNA damage sensitisation. Wild type and FUS-KO HeLa and SH-SY5Y cells were compared for their sensitivity to the DNA damage induced by camptothecin (CPT), an inhibitor of DNA topoisomerase I, and by etoposide (ETO), a topoisomerase II inhibitor. The viability after the genotoxic treatment was assessed by Trypan blue staining and counting of viable cells. Exposure of both HeLa FUS-KO and SH-SY5Y FUS-KO cells to either CPT or ETO resulted in a significantly stronger reduction in viability relative to their respective wild type cells (Figure 1F and Suppl. Figure 1F, respectively). Overall, these observations indicate that FUS is required for the maintenance of genome integrity.

### FUS knockout affects ATM-dependent signalling and recruitment of DDR factors

Next, we asked whether FUS could be directly involved in DDR signalling. HeLa wild type and FUS-KO cells were treated with ETO for 1 h and allowed to recover in ETO-free medium for 2 h (ETO release). Western blot analysis of cell extracts showed an increased phosphorylation of ATM, CHK2, BRCA1, and TRIM28, in addition to increased levels of γH2AX upon ETO treatment in both cell lines. Interestingly, 2 h after ETO release, while phosphorylation of most proteins started to reduce in WT cells, it remained higher in FUS-KO cells (Figure 2A). This indicates that the activation of the DDR occurred normally in the absence of FUS, but that DNA damage signalling persisted longer after the release from the genotoxic treatment. These data reinforce the concept either that cells lacking FUS generate more DNA damage than WT cells upon ETO treatment, or they repair the DNA damage less efficiently, or both.

**Figure 2.**
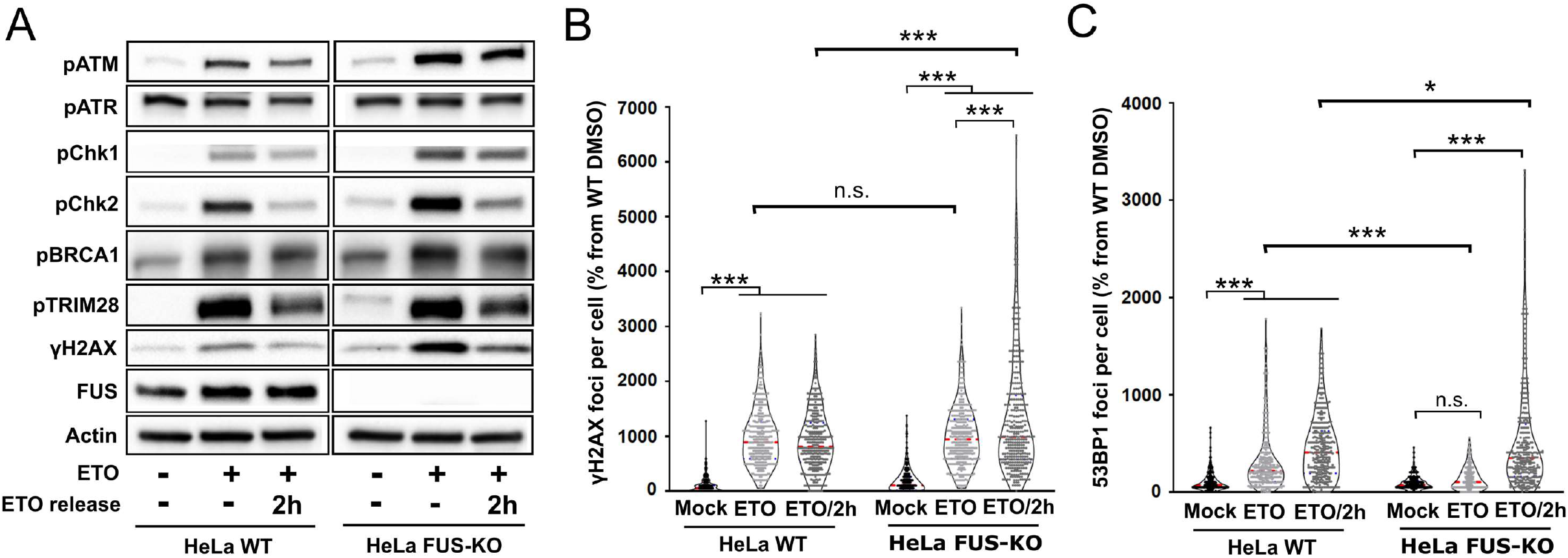
Loss of FUS perturbs DDR signalling and foci formation upon genotoxic insult. A. DDR activation in HeLa wild type and FUS-KO cells upon ETO treatment. Cells were treated with 10 µM ETO for 1 h and were allowed to recover in ETO-free media for 2 h (ETO release). Cells were collected at the indicated time points, lysed in the presence of phosphatase inhibitors, and processed for Western blotting. Actin was used as loading control. B. Quantification of ETO-induced γH2AX foci. Statistics: two-way ANOVA followed by Bonferroni post-hoc test. n.s., non-significant, *, p < 0.05, ***, p < 0.001. C. Quantification of ETO-induced 53BP1 foci analysed as in B.

To evaluate the impact of FUS depletion on DSB repair we used U2OS cells stably transfected with GFP reporter constructs that allow the measurement of HR and NHEJ-mediated repair (Gunn and Stark, 2012). The constructs are based on an engineered GFP gene containing recognition sites for the I-SceI endonuclease for enzymatic induction of DSBs. The starting constructs are GFP negative as the GFP gene is inactivated by an additional exon, or by mutations. Successful repair of the I-SceI-induced breaks by NHEJ or HR restores the functional GFP gene. The number of GFP positive cells, as counted by flow cytometry, provides a quantitative measure of the NHEJ or HR efficiency. Consistent with previous reports (Mastrocola et al., 2013; Wang et al., 2013) we observed that depletion of FUS affected both DSB repair pathways (Suppl. Figure 2A and 2B).

Spatiotemporal dynamics of DSB repair can be assessed in situ by monitoring the accumulation of DNA damage response proteins such as γH2AX, and 53BP1 at DSBs in microscopically visible, subnuclear foci. We thus determined whether the absence of FUS affected the formation of DNA damage foci. As shown in Figure 2B, after 1 h ETO exposure we observed a similar number of γH2AX foci in HeLa wild type and FUS-KO cells, indicating that the absence of FUS does not prevent the initial sensing of DSBs. However, after 2 h recovery from ETO treatment (ETO/2h), FUS-KO cells had significantly more γH2AX foci than wild type (Figure 2B and Suppl. Figure 2C). In addition, FUS-KO cells also showed significantly less 53BP1 foci than wild type cells after 1 h ETO (Figure 2C and Suppl. Figure 2D). Interestingly, 53BP1 foci formation was restored 2 h after recovery (Figure 2C). These results indicate that the absence of FUS results in a delay in the assembly of DNA repair complexes.

To gain insight in the molecular function of FUS during the early DDR events, we analysed the recruitment kinetics to DNA damage sites of two apical DDR factors, the 80 kDa subunit of the KU heterodimer, and NBS1, one of the subunits of the MRN. HeLa WT and FUS-KO cells transiently expressing GFP-KU80 or GFP-NBS1 were subjected to microirradiation with a 405 nm laser to induce a time-specific and localised DNA damage. Real time recording revealed that, in wild type cells, KU80 reached a peak of recruitment within 5 s after microirradiation, remaining there until the end of the assessment (180 s, Figure 3A and 3B). In FUS-KO cells, KU80 showed similar recruitment kinetics but its accumulation was severely impaired. In contrast to KU80, NBS1 showed slower recruitment kinetics, reaching a peak after about 130 s from microirradiation (Figure 3C and 3D). However, in the absence of FUS, NBS1 was more efficiently recruited compared to wild type cells. Transfection levels of GFP-tagged proteins in WT and FUS-KO cells were comparable as analysed by the raw fluorescence intensity (Suppl. Figure 3A and 3B). Overall, these observations demonstrate that FUS plays an apical role in DDR activation, in particular in the retention of KU80 at the DNA broken ends. In addition, our results suggest that, when stable KU80 binding is impaired, the MRN complex can more effectively gain access to the DNA ends.

**Figure 3.**
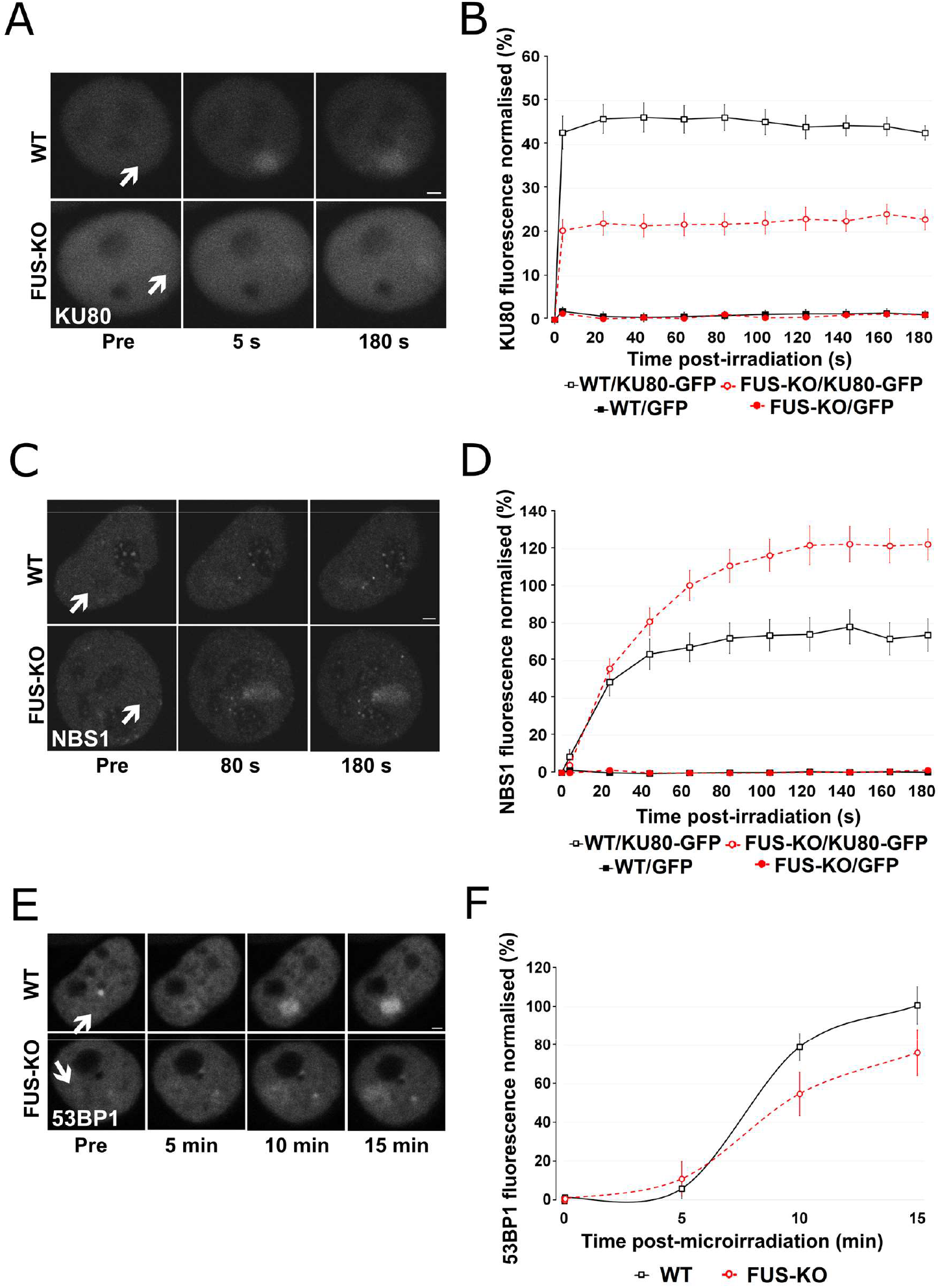
Loss of FUS changes the pattern of recruitment of HR and NHEJ-related proteins to DSBs. A. HeLa WT and HeLa FUS-KO cells were transiently transfected with KU80-GFP expressing plasmids. Images of selected time points, arrows indicate microirradiated area. Scale bar: 2 µm. B. Time course of the normalised fluorescence intensity of KU80-GFP protein at the microirradiated sites, average ± SEM for 10 cells in each experiment, performed twice (total of 20 cells per group). C. HeLa WT and HeLa FUS-KO cells were transiently transfected with a NBS1-GFP expressing plasmid. Images of selected time points, arrows indicate microirradiated area. Scale bar: 2 µm. D. Time course for NBS1-GFP analysed as in B. E. HeLa WT and FUS-KO cells were transiently transfected with 53BP1-GFP expressing plasmids. Images of selected time points, arrows indicate microirradiated area. Scale bar: 2 µm. F. Time course for 53BP1-GFP analysed as in B.

We next studied the recruitment of the effector protein 53BP1 in HeLa wild type and FUS-KO cells (Figure 3C). Consistent with the delayed appearance of 53BP1 foci observed in ETO-treated FUS-KO cells, we observed a reduction in the recruitment of GFP-53BP1 in the first 15 min after microirradiation in the absence of FUS (Figure 3E and 3F).

### FUS promotes the recruitment of SFPQ to sites of DNA damage

We reported earlier the interactome analysis of FUS (Reber et al., 2016). Gene Ontology (GO) analysis of this dataset revealed a substantial number of proteins that are involved in the DDR, including PARP1/2, XRCC6/KU70 and XRCC5/KU80 (Table 1). In addition, we also found a significant enrichment for RNA-binding proteins with low complexity domains (LCDs, Table 2), some of which have already been linked to DNA damage repair. Among potential FUS interactors, we further explored the functional interaction with SFPQ, because of its involvement in the DDR (Jaafar et al., 2017; Salton et al., 2010) and because, together with FUS, it is an essential structural component of paraspeckles (Hennig et al., 2015). Paraspeckles are nuclear membrane-less compartments that are built on the interaction of FUS and SFPQ with the NEAT1 lncRNA (Hennig et al., 2015; Naganuma et al., 2012) and whose integrity depends on liquid-liquid phase separation (LLPS, (Yamazaki et al., 2018)).

**Table 1.** DDR-related FUS interactors. This table was compiled from (Reber et al., 2016).

| Uniprot IDs | Protein Names |
| --- | --- |
| **Q92499** | ATP-dependent RNA helicase DDX |
| **P54132** | Bloom syndrome protein |
| **P49674** | Casein kinase I isoform epsilon |
| **P24941** | Cyclin-dependent kinase 2 |
| **Q03468** | DNA excision repair protein ERCC-6 |
| **P11388** | DNA topoisomerase 2-alpha |
| **Q92547** | DNA topoisomerase 2-binding protein 1 |
| **O60870** | DNA/RNA-binding protein KIN17 |
| **P29372** | DNA-3-methyladenine glycosylase |
| **P78527** | DNA-dependent protein kinase catalytic subunit |
| **P24928** | DNA-directed RNA polymerase II subunit RPB1 |
| **Q9Y5B9** | FACT complex subunit SPT16 |
| **Q08945** | FACT complex subunit SSRP1 |
| **Q9UBU8** | Mortality factor 4-like protein 1 |
| **Q15014** | Mortality factor 4-like protein 2 |
| **Q15233** | Non-POU domain-containing octamer-binding protein |
| **P06748** | Nucleophosmin |
| **P09874** | Poly [ADP-ribose] polymerase 1 |
| **Q9UGN5** | Poly [ADP-ribose] polymerase 2 |
| **Q9C0J8** | pre-mRNA 3' end processing protein WDR33 |
| **Q9UMS4** | Pre-mRNA-processing factor 19 |
| **Q9HCS7** | Pre-mRNA-splicing factor SYF1 |
| **P25789** | Proteasome subunit alpha type-4 |
| **P35251** | Replication factor C subunit 1 |
| **P35250** | Replication factor C subunit 2 |
| **P40938** | Replication factor C subunit 3 |
| **P35249** | Replication factor C subunit 4 |
| **P40937** | Replication factor C subunit 5 |
| **Q96PK6** | RNA-binding protein 14 |
| **Q9Y230** | RuvB-like 2 |
| **P23246** | Splicing factor, proline- and glutamine-rich |
| **Q9NTJ3** | Structural maintenance of chromosomes protein 4 |
| **Q13263** | Transcription intermediary factor 1-beta |
| **P07948** | Tyrosine-protein kinase Lyn |
| **Q14694** | Ubiquitin carboxyl-terminal hydrolase 10 |
| **P13010** | X-ray repair cross-complementing protein 5 |
| **P12956** | X-ray repair cross-complementing protein 6 |

**Table 2.** LCD-containing FUS interactors. This table was compiled from Hennig *et al*. JBC 2015 (Hennig et al., 2015) and Couthouis et al PNAS 2011 (Couthouis et al., 2011).

| Gene Name | Uniprot | Repeats in PLD | DDR involvement |
| --- | --- | --- | --- |
| **HNRNPA1** | P09651 | 2 x [G/S]Y[G/S] |  |
| **HNRNPA2/B1** | P22626 | 7 x [G/S]Y[G/S] |  |
| **HNRNPF** | P52597 | 2 x YXXQ, 5 x [S/G]Y[S/G] |  |
| **HNRNPH1** | P31943 | 7 x [G/S]Y[G/S] |  |
| **HNRNPH3** | P31942 | 6 x [G/S]Y[G/S] |  |
| **HNRNPK** | P61978 | 2 x [G/S]Y[G/S] | (Moumen et al., 2013) |
| **HNRNPR** | O43390 | 1 x YNQ, 1 x YGQQ | (Sui et al., 2015) |
| **HNRNPUL1** | Q9BUJ2 | 8 x YXQ, 8 x [G/S]Y[G/S] | (Harrison and Shorter, 2017; Polo et al., 2012) |
| **MATR3** | P43243 |  | (Salton et al., 2010) |
| **RBM14** | Q96PK6 | 19 x Y[G/N/A/S]AQ, 2 x | (Simon et al., 2017) |
| **SFPQ** | P23246 | 1 X SYQ | (Rajesh et al., 2011; Salton et al., 2010) |
| **TAF15** | Q86X94 | 12 x [G/S]Y[G/S], 9 x Y[G/S]Q |  |

To determine whether SFPQ function in DNA repair is dependent on FUS, we first characterised the recruitment kinetics of these two proteins in HeLa cells (Figure 4). HeLa WT cells were transiently transfected with a plasmid expressing GFP-tagged FUS or SFPQ and were then subjected to laser microirradiation to induce a time-specific and localised DNA damage, as confirmed by anti-γH2AX staining (Figure 4A). Upon microirradiation, while FUS is promptly recruited and reaches a maximum after 40 s, the redistribution of SFPQ to laser-induced damage sites was slower reaching a peak after approximately 100 s (Figure 4B and 4C). Then, we examined whether the recruitment of SFPQ is altered in the absence of FUS. As shown in Figure 4D and 4E, SFPQ accumulation was severely delayed and impaired in FUS-KO cells, indicating that the recruitment of SFPQ to DSB sites is FUS-dependent (Figure 4D and E, and Suppl. Video 1). Because laser microirradiation also generates SSBs, we tested whether the lack of FUS also affected the recruitment of XRCC1, a protein involved in base excision repair and the single-strand break repair pathway, by co-transfecting FUS-KO cells with both GFP-SFPQ and RFP-XRCC1. XRCC1 was efficiently recruited to DNA damage sites after laser microirradiation not only in WT cells but also in FUS-KO cells indicating that FUS is specifically required for the recruitment of SFPQ (Suppl. Figure 4).

**Figure 4.**
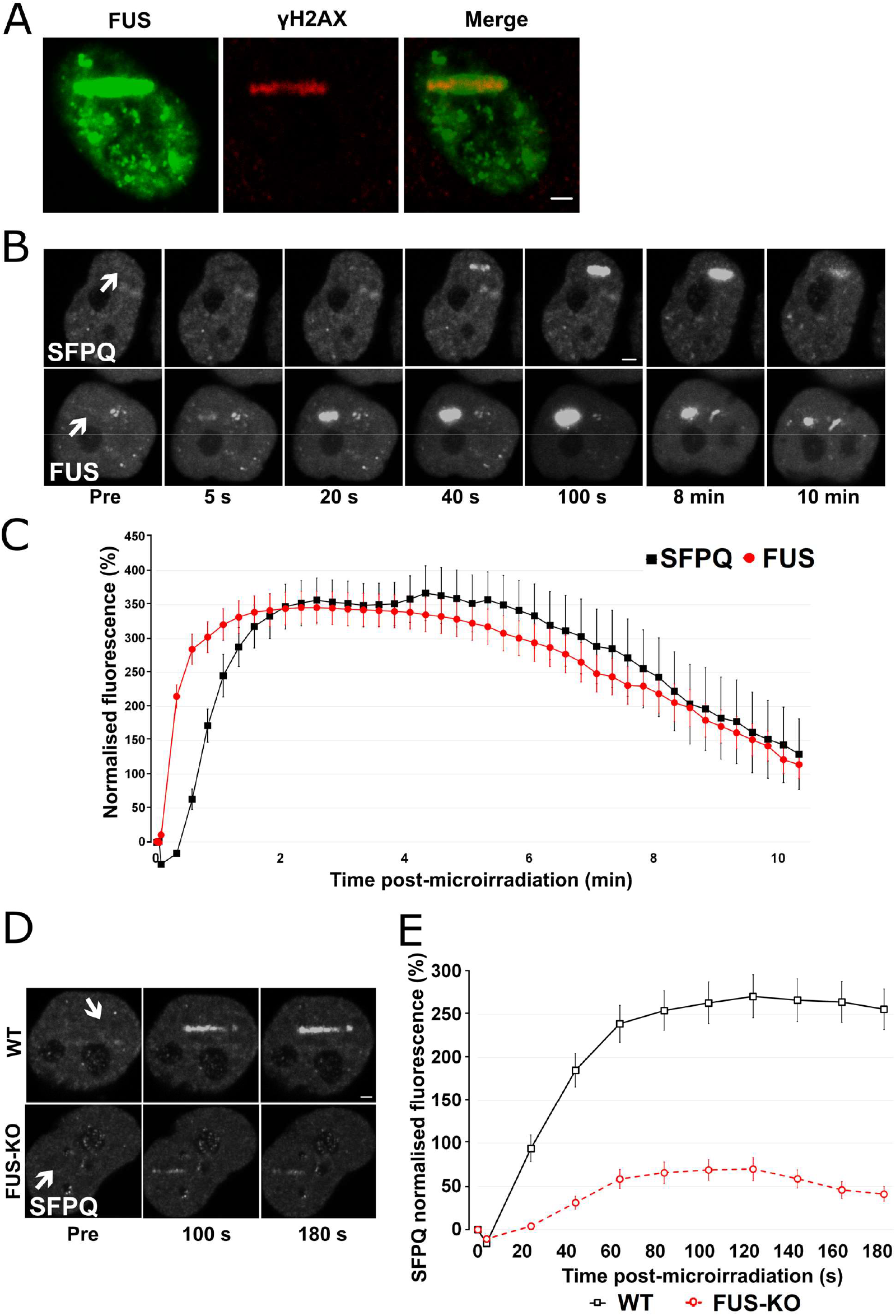
FUS recruitment to DSBs precedes SFPQ and its absence strongly reduces SFPQ accumulation. A. HeLa cells were transiently transfected with a GFP-tagged-FUS expression plasmid and then laser microirradiated. Cells were then immunostained for γH2AX. H2AX is phosphorylated at laser microirradiation sites and is co-localised with GFP-FUS. Scale bar: 2 µm. B. WT HeLa cells were transiently transfected either with GFP-SFPQ or GFP-FUS plasmid and and then laser microirradiated. Scale bar: 2 µm. C. Comparison of FUS and SFPQ recruitment kinetics. Data are plotted as normalised average ± SEM. D. HeLa WT and HeLa FUS-KO cells were transiently transfected with a GFP-SFPQ expressing plasmid and laser microirradiated. Scale bar: 2 µm. See also Suppl. Video 1. E. Quantification of SFPQ recruitment in WT vs FUS-KO cells. In all graphs, data are plotted as normalised average ± SEM.

Both FUS and SFPQ contain low complexity domains (LCDs) that were shown to liquid-liquid phase separate and to form hydrogels *in vitro* (Lee et al., 2015; Yamazaki et al., 2018). FUS is able to drive LLPS at low protein concentrations and physiological salt concentrations (Wang et al., 2018). Thus, we next tested whether the recruitment of FUS and SFPQ at DNA damage sites is dependent on LLPS and, more specifically, whether the recruitment of SFPQ is dependent on FUS-induced LLPS. We performed microirradiation experiments in the presence of chemicals that were previously shown to disrupt phase separation *in vivo*. First, we exposed HeLa cells to 1,6-hexanediol (1,6-HD), an aliphatic alcohol that is known to dissolve various cytoplasmic and nuclear membrane-less compartments *in vivo* by disrupting their multivalent hydrophobic interactions (Allodi et al., 2016; Kroschwald et al., 2015; Updike et al., 2011; Yamazaki et al., 2018), and which was shown to partially dissolve FUS polymers *in vitro* (Allodi et al., 2016; Kato and McKnight, 2018). We incubated HeLa cells transiently transfected with GFP-FUS for 30 minutes with 2% 1,6-HD prior to laser microirradiation. At this concentration, we observed that cells were able to recover their normal morphology 2 h after withdrawal from the alcohol (Supp. Figure 5A). Cajal Bodies were partially disrupted, while nuclear speckles were affected to a lesser extent (as shown by anti-coilin and anti-SC35 staining, Supp. Figure 5B and 5C). These observations indicate that this 1,6-HD treatment condition was mild enough not to disrupt all subcellular structures. Upon incubation with 1,6-HD, we observed a reduction in FUS relocation to DA damage sites (Figure 5A and Suppl. Figure 6A), an effect that was even more dramatic on the recruitment for SFPQ (Figure 5B and Suppl. Figure 6B). As control, we performed experiments in the presence of 2% 2,5 hexanediol (2,5-HD), a less hydrophobic isomer of 1,6-HD that does not affect LLPS (Allodi et al., 2016; Kato and McKnight, 2018). The presence of 2,5-HD did not affect FUS or SFPQ relocation (Figure 5A and 5B, and Suppl. Figure 6A and 6B).

**Figure 5.**
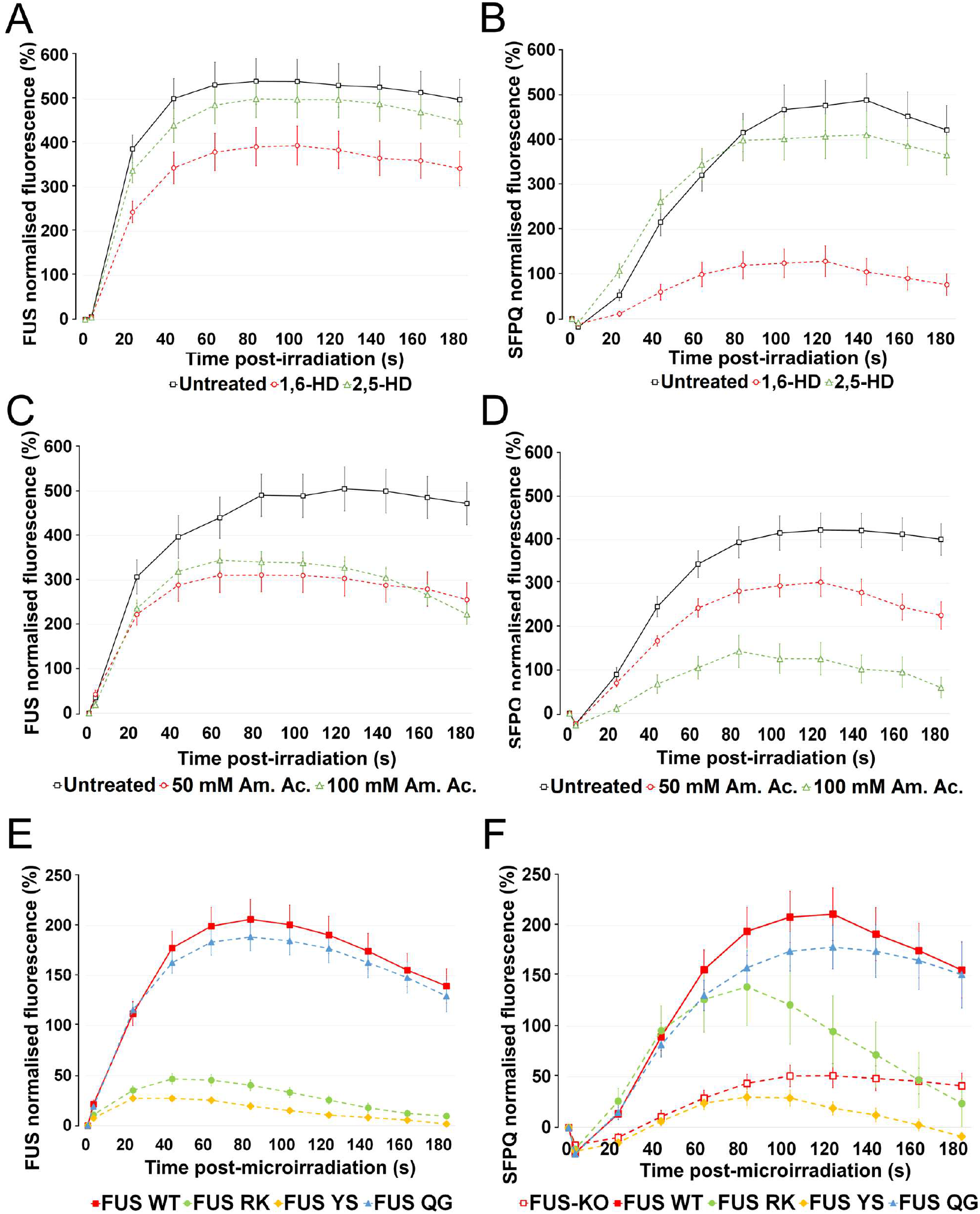
Liquid-liquid phase separation (LLPS) and LLPS-competent FUS are required for SFPQ recruitment to DNA damage sites. A. HeLa WT cells were transiently transfected with a GFP-FUS expressing plasmid and incubated with either 2% 1,6-HD or 2% 2,5-HD for 30 minutes prior to laser microirradiation. B. HeLa WT cells were transiently transfected with a GFP-SFPQ expressing plasmid and treated as in A. C. HeLa WT cells were transiently transfected with a GFP-FUS expressing plasmid and incubated with 50 mM or 100 mM ammonium acetate for 30 minutes prior to laser microirradiation. D. HeLa WT cells were transiently transfected with a GFP-SFPQ expressing plasmid and treated as in C. E. HeLa FUS-KO cells were transiently transfected with a mCherry FUS construct (WT or the mutants RK, YS or QG) and submitted to laser microirradiation. F. Recruitment of SFPQ-GFP in FUS-KO cells either not transfected with FUS or transfected with FUS WT, FUS RK, FUS YS or FUS QG.

**Figure 6.**
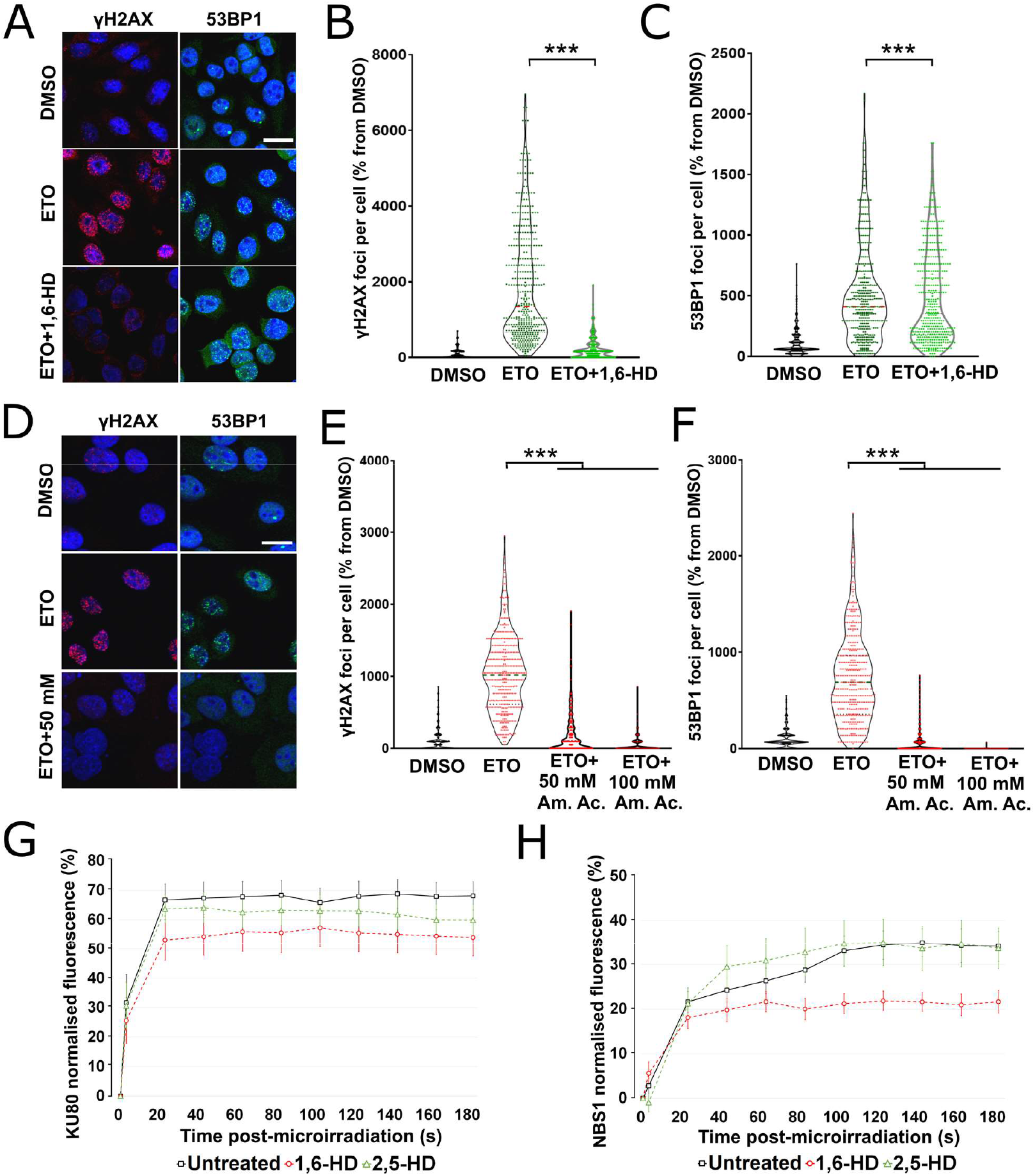
LLPS is required for γH2AX and 53BP1 foci formation and proper recruitment of HR and NHEJ-related proteins. A. Representative confocal micrographs of HeLa cells that were treated with ETO alone or with ETO and 1,6-HD prior to immunostaining for γH2AX or 53BP1. Scale bar, 20 µm. B. Quantification of γH2AX foci in the experiment in panel A. 200 cells were analysed per condition and experiments were performed in duplicate. Statistics: by one-way ANOVA followed by Bonferroni post-hoc test (*** p < 0.001). C. Quantification of 53BP1 foci in the experiment in panel A. Statistics as in Panel B. D. Representative confocal micrographs of HeLa cells that were treated with ETO alone or with ETO and 50 mM or 100 mM ammonium acetate for 30 min prior to immunostaining for γH2AX and 53BP1. Scale bar represents 20 µm. E. Quantification of γH2AX foci in the experiment in panel D. Statistics as in Panel B. F. Quantification of 53BP1 foci in the experiment in panel D. Statistics as in Panel B. G. HeLa WT cells were transiently transfected with GFP-KU80 expressing plasmids and then incubated with either 2% 1,6-HD or 2% 2,5-HD for 30 min prior to laser microirradiation. H. HeLa WT cells were transiently transfected with GFP-NBS1 expressing plasmids and then incubated with either 2% 1,6-HD or 2% 2,5-HD for 30 min prior to laser microirradiation.

To further substantiate the role of LLPS in FUS and SFPQ accumulation at laser-induced DNA damage sites we assessed the effect of ammonium acetate (Am. Ac.), which promptly permeates cells and inhibit RNA-protein gelation without perturbing intracellular pH (Hamaguchi et al., 1997; Jain and Vale, 2017). Therefore we incubated cells for 30 min prior to laser microirradiation with either 50 mM or with 100 mM Am. Ac.. These concentration of Am. Ac. had been shown to lead to the disappearance of RNA clustering and nuclear speckles within minutes (Jain and Vale, 2017), and to impair DNA damage repair (Pessina et al., 2019). As shown in Figure 5C and Suppl. Figure 6C, similarly to what we had observed in the presence of 1,6-HD, Am. Ac. severely affected the relocation of both FUS and SFPQ to DNA damage sites (Figure 5D and Suppl. Figure 6D). Overall, these results support the idea that LLPS occurs at sites of DNA damage and is required for the efficient recruitment of FUS and SFPQ.

We then hypothesised that the observed impairment in SFPQ recruitment observed in FUS-KO cells might be due to the absence of FUS-induced LLPS. To test this hypothesis, we took advantage of LLPS-deficient FUS constructs characterised by Wang and colleagues (Wang et al., 2018). We tested the FUS PLD Y → S (Y/S) and RNA-binding domain R → K (R/K) variants which strongly affect phase separation and, as a control, the FUS PLD Q → G (Q/G) variant which instead affects the hardening of droplets (Wang et al., 2018). These FUS constructs were transiently transfected as mCherry fusions in HeLa FUS-KO cells. As shown in Figure 5E, while wild type FUS and the Q/G FUS variant were similarly recruited to DNA damage sites, the accumulation of Y/S and R/K FUS variants was significantly reduced (Figure 5E and Suppl. Figure 7). To test whether the expression of these FUS variants influence the recruitment of SFPQ, HeLa FUS-KO cells were transiently co-transfected with plasmids expressing GFP-SFPQ and one of the mCherry-FUS proteins (WT, or Y/S, or Q/G, or R/K). As shown in Figure 5F, while complementation of FUS-KO cells with the WT FUS or the Q/G FUS variant was able to increase SFPQ recruitment, expression of the Y/S FUS variant did not improve SFPQ relocation to laser-induced DNA damage sites. Interestingly, in the presence of the R/K FUS variant, SFPQ was initially recruited but was then prematurely released from the DSB sites (Figure 5F and Suppl. Figure 7). Overall, these observations implicate a requirement for FUS-driven LLPS for the efficient recruitment and retention of SFPQ at DNA damage sites.

**Figure 7.**
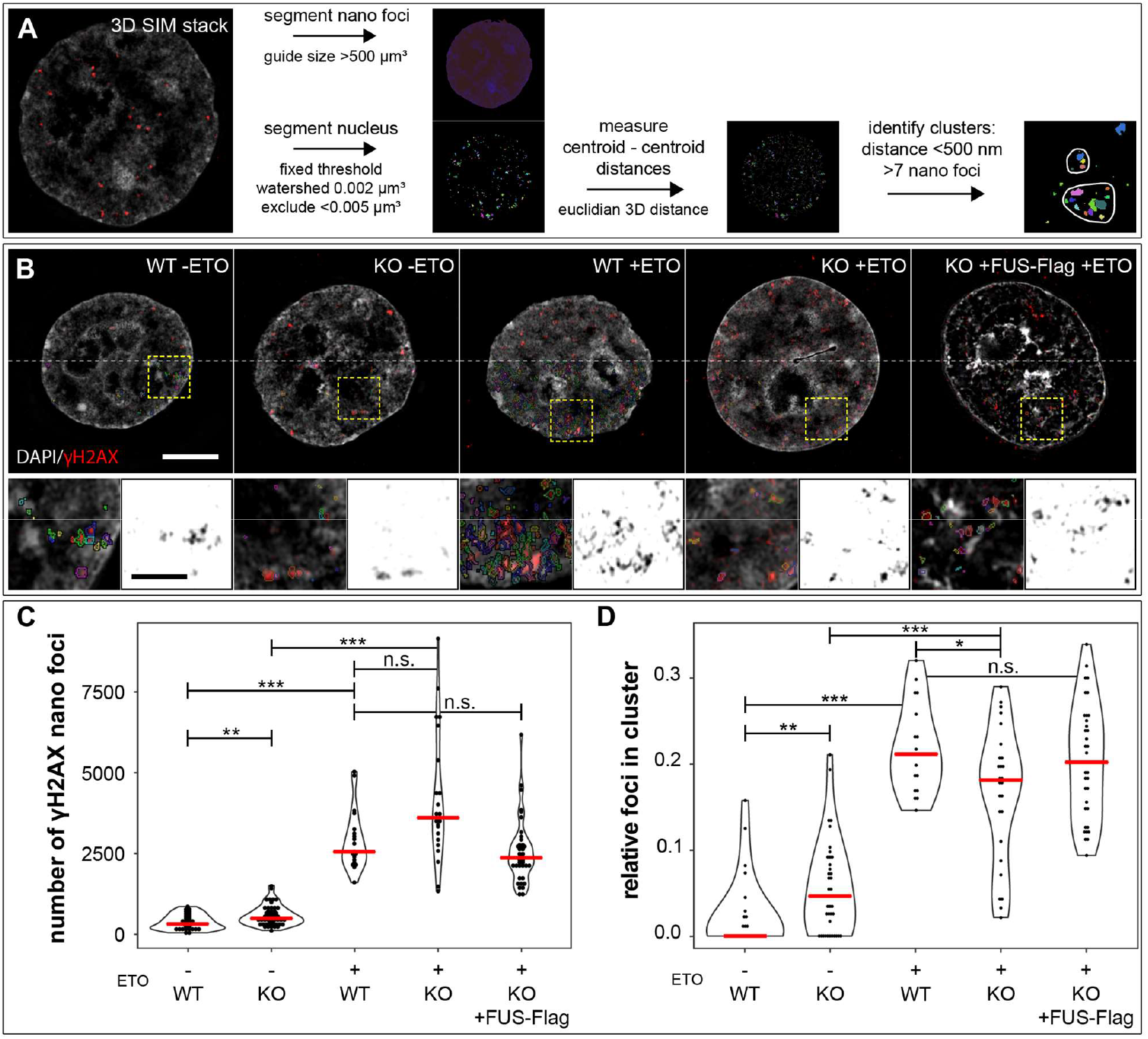
FUS is required for γH2AX nano-foci clustering. A. Image analysis workflow for 3D-SIM γH2AX cluster analysis. First nuclei and γH2AX nano-foci are segmented with the indicated constrains. Then the centroid-centroid distances for the nano-foci are computed and nano-foci with a centroid-centroid distance smaller than 500 nm are assigned to the same cluster. Clusters are quantified as structures that contain more than 7 nano-foci. B. Images of HeLa wild type and FUS-KO cells treated for 2 hours with etoposide or DMSO, respectively. Additionally, HeLa-FUS-KO cells transiently transfected with FUS-Flag plasmid were analysed after etoposide treatment. The main images show the merge of DNA (DAPI) and γH2AX nano-foci in pseudo colour. The lower half of the image shows the outline of the segmented nano foci. Crop areas are highlighted with yellow boxes and the magnified regions are shown below, again as merge image and as inverted grey scale image for the γH2AX channel only. C. Quantification of nano-foci per cell in the different conditions indicated. Red lines indicate the median. D. Frequency of nano-foci found in clusters, as compared to the total number of nano-foci in the cell. Red lines indicate the median. Statistics: Wilcoxon’s Rank sum Test. *, p < 0.05, **, p < 0.01 and ***, p < 0.001.

### Liquid-liquid phase separation is required to maintain the integrity of DNA damage foci

We then speculated that the actual formation of DDR foci might involve some form of phase separation. We first tested this hypothesis by exposing HeLa cells to either 2% 1,6-HD, or to 50 mM or 100 mM ammonium acetate, together with ETO for 30 min. Then, cells were fixed and stained with γH2AX or 53BP1 antibodies to assess DDR foci formation. 1,6-HD treatment severely impaired the formation of γH2AX (Figure 6A and 6B). In contrast, 53BP1 foci were only partially dissolved (Figure 6A and 6C). In the presence of Am. Ac., γH2AX foci formation was severely impaired similarly to what we had observed with the aliphatic alcohol (Figure 4D and 6E). Am. Ac. also affected 53BP1 foci formation (Figure 6D and 6F), as previously reported (Pessina et al., 2019). In addition to compromising foci integrity, 1,6-HD and Am. Ac. treatment also compromised the phosphorylation of ATM, H2AX and TRIM28 (Suppl. Figure 8A). To test whether the recruitment of key DDR factors to sites of DNA damage is also dependent on LLPS, we assessed the effect of the aliphatic alcohols on the recruitment of GFP-KU80 (Figure 6G and Suppl. Figure 8B) and GFP-NBS1 (Figure 6H and Suppl. Figure 8C). We observed a reduction in the recruitment of both DDR factors in the presence of 1,6-HD, but not in the presence of the control alcohol 2,5-HD. Overall, these data indicate that LLPS is required for the proper formation of DNA damage foci and the activation of the DDR signalling cascade.

### FUS is required for γH2AX clusters formation

We have recently shown using structured illumination microscopy (3D-SIM) that γH2AX foci consist of spatially clustered γH2AX nano-foci (Natale et al., 2017). Given that FUS-KO cells have defective recruitment of DNA damage factors and formation of DNA damage foci, we next assessed whether FUS may be required for the proper organisation of the three-dimensional (3D) arrangement of γH2AX-decorated chromatin. We thus quantified γH2AX nano-foci in WT and FUS-KO cells treated with 10 µM ETO for 1 h using 3D-SIM.

Figure 7A displays the workflow for foci segmentation and cluster identification. Consistent with the data obtained by confocal microscopy (Figure 1D), DMSO-treated WT control cells exhibited a low number of γH2AX nano-foci, while foci were significantly increased in control FUS-KO cells (Figure 7B and 7C). Upon ETO treatment, we did not observe a significant difference between the total number of nano-foci in WT versus FUS-KO cells (Figure 7C); similar to the results we obtained by confocal microscopy.

We then studied the spatial distribution of the induced γH2AX nano-foci in terms of cluster formation. To address this, we segmented all nano-foci in the nucleus and calculated the Euclidean distances between the centroids of each nano-focus. Nano-foci with a centroid-centroid distance smaller than 500 nm were assigned to the same nano-foci cluster. We then counted the nano-foci clusters that contained a minimum of 7 nano-foci and calculated the fraction of γH2AX nano-foci contained in clusters compared to the total number of γH2AX nano-foci per cell. Unsurprisingly, considering the extremely low number of γH2AX nano-foci in DMSO-treated WT cells, we found the fraction of γH2AX nano-foci clusters in these cells to be extremely low. In contrast, the fraction of clusters was higher in DMSO treated FUS-KO cells, probably due to the increased total number of γH2AX nano-foci in FUS-KO cells. Upon ETO treatment, while WT cells showed a significantly increased fraction of γH2AX nano-foci organised in typical clusters, these were significantly reduced in FUS-KO cells (Figure 7B and 7D), indicating that FUS is required for the proper organisation of the 3D arrangement of γH2AX-decorated chromatin. To support the conclusion, we transiently transfected HeLa FUS-KO cells with a FUS-Flag construct and quantified the number of γH2AX nano-foci and clusters upon ETO treatment (Figure 7B, right panel). Differently from untransfected FUS-KO cells, FUS-Flag transfected FUS-KO cells showed a similar total number of total clustered nano-foci as WT cells (Figure 7C and 7D). These findings demonstrate that FUS is an additional factor, besides CTCF and direct chromatin contacts (Collins et al., 2020; Hwang et al., 2019; Natale et al., 2017), which is required for the orchestrated formation of γH2AX-decorated chromatin domains. These domains are the essential basis for the subsequent steps of DSB repair, and our results indicate that the defective DNA damage repair in FUS-KO cells is associated to inefficient clustering of γH2AX nano-foci.

## Discussion

The results reported here provide the first evidence that FUS plays an important early role in the DDR by promoting LLPS at DNA damage sites thereby physically contributing to the efficient recruitment of key DDR factors.

Previous reports have already implicated FUS in DNA damage repair (Mastrocola et al., 2013; Singatulina et al., 2019; Wang et al., 2013), and more recently review articles have proposed that FUS, and more generally RNA-binding proteins, contribute to the DDR and to DNA repair by promoting phase separation at damage sites (D’Alessandro and d’Adda di Fagagna, 2017; Kai, 2016). However, *in vivo* experimental evidence of their molecular function in this process remained to be established.

We screened DDR proteins for their recruitment to laser-induced damage sites in FUS knock-out cell lines. We found that FUS is required at a very early step for the retention at DSBs of the 80 kDa subunit of the DSB sensor KU70/80. Interestingly, whereas in FUS-KO cells the retention of KU80 is impaired, the recruitment of NBS1, one of the three subunits of the MRN complex, is increased by more than 50%. These observations are consistent with the idea that, due to its high abundance and strong affinity for DNA, KU heterodimer is the first DDR factor that binds to the DNA broken ends irrespective of the cell cycle phase (Shibata et al., 2018). KU70/80 is then removed by the endonucleolytic activity of the MRN complex (Chanut et al., 2016; Myler et al., 2017) or by phosphorylation of KU70 (Lee et al., 2016). Therefore, KU70/80 is recruited but not efficiently retained in the absence of FUS allowing the MRN complex to successfully compete for the binding to the DNA broken ends. However, this is not sufficient for efficient repair, since we and others have demonstrated that the silencing of FUS affects both HR and NHEJ-mediated repair (Mastrocola et al., 2013; Wang et al., 2013). This justified the continuing quest for FUS-dependent events during the DDR.

Indeed, FUS is also required for the relocation of 53BP1 to DNA damage sites and consistently the formation of 53BP1 foci is delayed in FUS-KO cells. We also discovered that FUS is necessary for the recruitment to damage sites of SFPQ, an RNA-binding protein that had already been implicated in DNA damage repair (Bladen et al., 2005; Jaafar et al., 2017; Rajesh et al., 2011; Simon et al., 2017). Intriguingly, FUS and SFPQ are both core components of paraspeckles, phase-separated membrane-less organelles that assemble on the NEAT1 lncRNA, which induces liquid– liquid-phase separation via interaction with SFPQ/NONO (Yamazaki et al., 2018). The purified LCD of FUS forms a hydrogel at high concentration *in vitro* and this region is essential for paraspeckle formation *in vivo* (Hennig et al., 2015). Based on these observations it was tempting to hypothesise that FUS and SFPQ play a similar role in the DDR, inducing LLPS at DSBs to promote the efficient assembly of DNA repair complexes. Indeed, we show that efficient relocation of SFPQ requires LLPS-competent FUS.

Consistent with a physical role of FUS and –inducing LLPS– in triggering the formation of DDR protein assemblies around DSBs, we found that γH2AX foci are sensitive to the treatment with two different LLPS inhibitors, indicating that LLPS occurs very early during the DDR. In addition, LLPS inhibition impairs the recruitment of FUS, SFPQ, KU80 and NBS1, and affects DDR signalling. In fact, FUS forms LLPS-specific interactions with additional DNA damage repair proteins (Reber et al., BiorXiv, https://doi.org/10.1101/806158). Our findings are consistent with two recent studies showing that LLPS is necessary for the subsequent recruitment of the downstream effector 53BP1 to DNA damage foci (Kilic et al., 2019; Pessina et al., 2019).

To support the idea that FUS-dependent LLPS at DSBs is required for the efficient assembly of DNA repair complexes, we investigated the structure of H2AX foci formed in the absence of FUS using superresolution microscopy. We reported previously that γH2AX foci formed following IR exposure consist of spatially clustered nano-foci of ∼200 nm diameters (Natale et al., 2017). Here we characterised γH2AX foci in WT and FUS-KO cells treated with ETO. We found that FUS contributes to the spatial clustering of γH2AX foci. The impairment of the structural organisations of γH2AX nano-foci in FUS-KO cells upon genotoxic damage is reminiscent of what we observed upon depletion of the chromatin architectural protein CTCF (Natale et al., 2017). Thus, we propose that FUS exerts an early role in DDR by promoting LLPS. This, and the presence of CTCF, contributes to the 3D organisation of chromatin around DSBs enabling the activation of an efficient DDR.

The question remains open as to what recruits FUS to broken DNA ends. In this regard, the formation of a complex between KU and FUS is supported by several recent reports (Abbasi and Schild-Poulter, 2018; Morchikh et al., 2017). Based on our data and on the study by Alexandrov and colleagues who measured, clustered, and modelled the kinetics of recruitment and dissociation of 70 DNA repair proteins to laser-induced DNA damage sites (Aleksandrov et al., 2018), we propose that FUS relocates to DSBs where it promotes LLPS together with SFPQ. This stabilises KU70/80 on the broken DNA ends and leads to the further recruitment of multiple proteins required for DSB repair. Intriguingly, several of these proteins are RBPs, which supports the idea that RNAs are involved in DSB repair (Michelini et al., 2017; Pessina et al., 2019). How FUS and other RBPs are recruited to broken ends remains to be established. Most likely, this occurs through binding to poly-ADP ribose chains that are deposited at sites of DNA damage by PARP1, which is among the first proteins that are recruited (have half-times between 1.8 and 3.7 s (Aleksandrov et al., 2018)). Both FUS and the SFPQ/NONO heterodimer have been shown to bind to PAR (Krietsch et al., 2012; Mastrocola et al., 2013). While further work is required to understand the role of all the different RBPs that have been involved in DNA repair, and also of RNA itself, our data will facilitate mechanistic investigations to uncover their precise activities.

## Acknowledgments

This work was partially supported by the Swiss National Fond Sinergia grant no. CRSII3_136222 (to O.M., S.M.L), the UK Dementia Research Institute (M.D.R) which receives its funding from UK DRI Ltd, funded by the UK Medical Research Council, Alzheimer’s Society and Alzheimer’s Research UK, and the NOMIS Foundation (M.D.R). We are grateful to J. Stark for the U2OS cell lines stably expressing HR and NHEJ repair reporters. We are in debt to J. Wang and S. Alberti for having shared their FUS constructs, to S. Cerritelli for having provided the pEGFP-M27-H1 plasmid, and to A. Zippo for having supplied vectors to perform lentiviral infections. NBS1-GFP was kindly provided by A.Nussenzweig. We are grateful to Dr. Marco E. Bianchi for critical reading of the manuscript.

## Authors Contribution

B.R.L. conceived and performed the experiments and the data analysis for Figs. 1, 2, 3, 4, 5, 6 and associated supplementary data, and participated in preparing figures and tables. S.C.L. conceived and performed the experiments and the data analysis for Figs. 1 and 2 and associated supplementary data, and participated in preparing figures and tables. M.A. performed the experiments in Fig. 1A, 2A, and associated supplementary data. A.M. and A.R. performed the experiments and data analysis for Fig. 7. F.C. performed the experiments in Fig. 1E and 2A. S.R. generated knockout lines and FUS constructs. A.E.R., O.M., and M.-D.R. critically revised the manuscript. H.L. and M.C.C. supervised the experiments in Fig. 8 and critically revised the manuscript. S.M.L.B. conceived the study and wrote the manuscript. All co-authors provided discussion and data interpretation and contributed to the final version of the manuscript.

## Methods

### Cell lines, cell culture and treatments

HEK293T cells stably expressing Flag-tagged FUS and HeLa FUS-KO are described in (Reber et al., 2016). U2OS cells stably expressing HR and NHEJ repair reporters were a kind gift from J.M. Stark and are described in (Gunn and Stark, 2012). HeLa and SH-SY-5Y FUS-KO cells were generated by CRISPR-trap (Reber et al., 2018). All cell lines were tested for mycoplasma contamination and were found to be negative. Cells were cultured in Dulbecco’s Modified Eagle’s Medium (DMEM High Glucose, Euroclone) supplemented with 10% Foetal Bovine Serum (EuroClone), 2 mM L-Glutamine (EuroClone) 100 IU/ml Penicillin (Euroclone) and 0.1 mg/ml Streptomycin (Euroclone). Cells were grown at 37°C and 5% of CO2 in a humidified incubator.

Etoposide (ETO) treatment, where not specified otherwise, was performed for 1 h with 10 μM diluted in DMSO (Enzo Lifesciences). Where specified, cells were treated with either 2% 1,6-Hexanediol (1,6-HD, diluted in growth medium, Sigma-Aldrich) or 2% 2,5-Hexanediol (2,5-HD, diluted in growth medium, Sigma Aldrich) or 50 or 100 mM ammonium acetate (diluted in water, Sigma Aldrich) for 30 minutes.

Plasmid DNA transfections were performed using Lipofectamine 2000 (Invitrogen), while siRNA transfections were done using Lipofectamine RNAiMAX (Invitrogen), according to the manufacturer’s instructions. The synthetic siRNAs used in this study were purchased from Riboxx Life Sciences GmbH (Germany).

### Trypan blue assay

HeLa WT and FUS-KO cells were seeded at a concentration of 2 × 10^3^ cells/well in 96-wells plates. Cells were allowed to attach overnight and were then treated with increasing concentrations of etoposide (ETO: DMSO, 0.5 or 1 µM) or camptothecin (CPT: DMSO, 0.1 or 0.5 µM) for 18 h. Cells were then detached with trypsin and diluted 1:2 in Trypan blue. Living cells were then visualised and counted with a hemocytometer using a bright field microscope. Number of cells is a percentage of their respective control (DMSO).

### DNA constructs

NBS1-GFP was kindly provided by Dr. Nussenzweig (Kruhlak et al., 2006). KU80-EGFP was purchased from AddGene (#46958). SFPQ-EGFP was generated by subcloning the SFPQ ORF obtained from the Myc-PSF-WT plasmid (AddGene #35183). The FUS-EGFP plasmid was generated by inserting the FUS ORF into pcDNA6F-EGFP. XRCC1-RFP plasmid (pc1156) was generated by subcloning the XRCC1 obtained from a pEGFP-XRCC1 (pc1152) into pmRFP from a pmRFP-Lig4 (pc1079) (Mortusewicz et al., 2006). FUS-mCherry plasmids (WT, RK, YS and QG) were kindly provided by Dr. S. Alberti (Wang et al., 2018). All oligonucleotides used in this study are listed in the Supplementary Table 1.

### Protein collection, quantification, and western blotting

Adherent cells were washed twice with PBS and then scraped and digested on ice for 30 min with a homemade RIPA (50 mM Tris-HCl pH 7.5, 15 mM NaCl, 1% NP-40, 0.5% sodium deoxycholate and 0.1% SDS, supplemented with Protease Inhibitors [Roche] and Phosphatase Inhibitors [Sigma-Aldrich]). Cell extracts were centrifuged at 16,000xg for 20 min at 4°C and only the soluble fraction (supernatant) was used. Proteins were quantified using a validated BCA (Bicinchoninic Protein Assay Kit, EuroClone) protocol. For the western blotting, the same amount of proteins was loaded, diluted in a homemade sample buffer (6X Laemmli sample buffer: 1 M Tris-HCl pH 6.8, 10% SDS, 40% Glycerol, 12% β-mercaptoethanol and bromophenol blue). Running buffer was prepared with tris-glycine 1X and SDS 0.1% using a homemade gel (7-12% bis-acrylamide, Bio-Rad), which ran at 100 V for approximately 2:30 h. After the run, the proteins were transferred to a nitrocellulose membrane (0.45 µm pore size, Amersham). Transfer was performed at 80 V at 4°C for 1:30 to 2:30 h using a 20% methanol transfer buffer. Membranes were stained with Ponceau (Sigma) for 8 minutes and images were acquired using a ChemiDoc (BIO-RAD). After removal of the Ponceau by washing once with TBS-T (TBS 1X, Tween 0.05%), membranes were blocked for 1 h with 5% milk or 4% BSA (for phosphorylated proteins), diluted in TBS-T. Membranes were then incubated overnight at 4°C with a primary antibody diluted in 5% milk or 4% BSA (a list of all used antibodies and dilutions is provided in Supplementary Table 2). The membrane was then washed three times with TBS-T for 5 minutes and incubated with a secondary antibody diluted in 5% milk for 1 h at room temperature. The membrane was finally washed three times with TBS-T for 10 minutes. The acquisition was made using Cyanagen ECL according to manufacturer’s protocol, exposing the membrane in the trans illuminator ChemiDoc (BIO-RAD). The quantification of the signal intensity performed using Image Lab 6.0.

### Immunofluorescence, confocal imaging, and quantification

Cells were fixed with PFA 4% (diluted in PBS, pH 7.4) at RT for 15 min and, after 3 washes with PBS, permeabilised with 0.25% Triton-X100 in PBS for 5 min. Permeabilised cells were blocked in blocking solution (20% FBS, 0.05% Tween in PBS) for 1 h at RT. Primary antibodies were diluted in wash buffer (0.2% BSA in PBS) according to Supplementary Table 4, and incubated for 1 h at RT. After 3 washes with wash buffer, cells were incubated with the respective secondary antibodies (diluted in wash buffer) for 1 h at RT. After 3 washes with wash buffer, cells were counterstained with DAPI (Sigma Aldrich, 1 µg/ml in PBS) diluted in PBS for 10 min. Coverslips were washed twice with PBS and once with water and mounted onto microscope slides using an antifade mounting medium (FluorSave, Calbiochem). All antibodies used in the present study are listed in Supplementary Table 2.

Cell imaging was performed using a confocal microscope ECLIPSE - Ti A1 (Nikon). Between 6-10 images were taken per coverslip (randomly), using the 60x oil objective. Quantification of DNA damage foci (γH2AX and 53BP1) and subnuclear bodies (Cajal bodies and nuclear speckles) was performed using the software Fiji ImageJ 2.0. For that end, a mask of the nucleus was made using the DAPI staining (filter Gaussian Blur radius 2 followed by adjusting the threshold and then analysing particles). Nuclear foci were measured using the function “Find Maxima” (by setting a noise tolerance and dividing the final image by 255). For baseline experiment (WT vs FUS-KO), data are shown as the average number of foci per nucleus. For all other experiments, data are normalised by the WT DMSO group (which was considered 100%). At least 100 cells were counted per experiment, which were done at least in duplicate.

### Laser microirradiation

HeLa WT and/or FUS-KO cells were transiently transfected with the indicated GFP or mCherry-tagged plasmids two days before the irradiation. The following day, transfected cells were plated onto 35 mm plates with a glass bottom and allowed to attach overnight. 30 min prior to irradiation, the cell medium was replaced by a phenol red-free medium, containing 0.5 µg/ml of Hoechst (Sigma-Aldrich). Three images were taken as baseline (pre-irradiation) and then cells were irradiated for 4 s using the 405 nm laser at 25% power. The fluorescence intensity of the irradiated area and of two other nuclear regions of interest (ROIs) and one background ROI were assessed. The protein recruitment to the irradiated area was analysed by removing the background fluorescence intensity from each of the other ROIs, then calculating the percentage change at each time point from the average of the three baseline images (pre-irradiation). Finally, the difference between the fluorescence signal in the irradiated region and the average of the two control regions in the nucleus was calculated. The formula used to calculate the protein recruitment is: Recruitment (%) = % from baseline in irradiated ROI – [(% from baseline in control ROI 1 + % from baseline in control ROI 2) / 2]. All the microirradiation experiments were performed using the confocal microscope ECLIPSE - Ti A1 (Nikon). Experiments were done in duplicate and 10 cells were assessed per experiment (to a total of 20 cells per group).

### HR and NHEJ repair reporter assays

DSB repair assays were performed in DR-GFP or EJ5-GFP U2OS cell lines (Gunn and Stark, 2012). Briefly, cells were transfected with siRNAs. Two days later cells were co-transfected with a plasmid expressing I-SceI (pCBA-I-SceI) together with indicated siRNAs using Lipofectamine 2000 (Invitrogen). Cells were harvested after three days post transfection and subjected to flow cytometric analysis to identify and quantify GFP-positive cells (BD FACS Calibur, CellQuest software). The repair efficiency was scored as the percentage of GFP-positive cells and data were normalised by a control siRNA treatment in each individual experiment. Experiments were done in triplicates and at least 10,000 cells were assessed per experiment.

### 3D SIM Imaging and analysis

Cells were plated on high precision coverslips (18×18 mm, #1.5 thickness with low variance). After treatment, cells were fixed with 2% formaldehyde at RT for 10 min and then washed 3 times (first time by removing only 2/3 of fixation solution) with 0.02% Tween 20 in PBS (PBST). Cells were permeabilised with 0.5% TritonX-100 in PBST for 10 min and blocked with 2% bovine serum albumin (BSA) in PBST for 1 h at RT. After blocking, primary antibody diluted in blocking solution (1:#### give clone name) was added for 1 h at RT, followed by 3 washes with PBST. Secondary antibody diluted in blocking solution was then added for 1 h at RT (1:#### give name and cataloge #), followed by 3 washes with PBST. Samples were postfixed with 4% formaldehyde for 10 min at RT and then washed 3 times with PBST. A DAPI counterstaining (Sigma Aldrich, 1 µg/ml in PBST) was done for 10 min at RT and then washed once with PBST and once with ddH2O before mounting onto a microscope slide using Vectashield antifade mounting medium and sealed with nail polish.

A two-step analysis of γH2AX clusters was done with Volocity software 6.1.2 (Perkin Elmer) and with the software “R” (R foundation). First, aligned 3D-SIM RGB image stacks were used in Volocity and the respective channels were separated for the segmentation of γH2AX structures and DAPI stained nuclei. Hereof, γH2AX segmentation was performed for all cells with the following commands: 1. “Find Objects” (Threshold using: Intensity, Lower: 32, Upper: 255), 2. “Separate Touching Objects” with an Object size guide of 0.002 µm^3^ and 3. “Exclude Objects by Size”, excluding structures < 0.005 µm^3^. Besides volume measurements, centroid position of each γH2AX volume was always registered. For the challenging segmentation of nuclei, we used the commands “Find Objects”, “Dilate”, “Erode” and “Fill Holes in Objects” with specific settings. To obtain only γH2AX structures within a nucleus, the “Intersect” and “Compartmentalize” command was used. In a second step, the provided information of γH2AX structures (centroid position etc.) was transferred to “R”. Here the euclidian centroid to centroid distances were computed for all nano foci structures within one nucleus and then nano foci with a centroid-centroid distance smaller than 500 nm were assigned to the same cluster. Obtained clusters were filtered for structures that contain a minimum of 7 nano foci.

### Statistical analysis

Bar graphs show average ± S.E.M., while boxplot represent median and quartiles (mean is represented by an x). Statistical analysis was performed using the software IBM SPSS Statistics Version 26. Comparison of two groups was done by Student’s t-test, while more groups where compared by one-way ANOVA. Two variable comparisons were performed using two-way ANOVA. All post-hoc analysis, when necessary, were done using Bonferroni test. Values of p<0.05 were considered significant. Significant values are shown by * p<0.05, ** p<0.001 and *** p<0.001.

