## Supplementary Material for "FUS-dependent liquid-liquid phase separation is an early event in double-strand break repair"

[§] Present address: Institute of Molecular Biology (IMB)¸ Ackermannweg 4, 55128 Mainz, Germany

Running Title: FUS-dependent LLPS in DDR signalling

* Corresponding author: Silvia M.L. Barabino

Department of Biotechnology and Biosciences

University of Milano-Bicocca, Piazza della Scienza, 2

I-20126 Milano, Italy

**SUPPLEMENTARY TABLES**

Supplementary Table 1. **Sequences of oligonucleotides and siRNAs, related to experimental procedures**.

| **Name** | **Sequence (5'-3')** |
|---|---|
| **siSFPQ** | GGCAAAGGAUUCGGAUUUAuu |
| **siNONO** | GGGGUGGUAUUAAACAAGUca |
| **siXRCC5** | AAGAGCUAAUCCUCAAGUCuu |
| **siTOPBP1** | CUCACCUUAUUGCAGGAGAdTdT |
| **siLIG4** | GGCAUCUGGUAAGCUCGCAUCUAAA |
| **siFUS1** | AGCCCAUGAUUAAUUUGUAtt |
| **siFUS_sh** | GGACAGCAGCAAAGCUAUAtt |
| **siRNA pool negative control** | Catalogue number: SKU# K-00100 (iBONI - Riboxx's design) |

Supplementary Table 2. **List of antibodies used in the study.**

| |
|---|
| ***Primary antibodies (Immunofluorescence)*** |
| Mouse α-γH2AX: Abcam ab26350, dilution 1:100 |
| Rabbit α-53BP1: NovusBio NB100-305, dilution 1:200 |
| Rabbit α-Coilin: A.I. Lamond, dilution 1:100 |
| Rabbit α-SC35, dilution 1:100 |
| ***Primary antibodies (Western Blot)*** |
| Rabbit α-γH2AX: Cell Signalling #9718, dilution 1:1000 |
| Rabbit α-pATM: Cell Signalling #5883, dilution 1:1000 |
| Rabbit α-pATR: Cell Signalling #2853, dilution 1:1000 |
| Rabbit α-pCHK1: Cell Signalling #2348, dilution 1:1000 |
| Rabbit α-pCHK2: Cell Signalling #2197, dilution 1:1000 |
| Rabbit α-pTRIM28: Bethyl A300-767A-T, dilution 1:1000 |
| Rabbit α-pBRCA1: Cell Signalling #9009, dilution 1:1000 |
| Rabbit α-FUS: Homemade, Dr. M.-D. Ruepp, dilution 1:3000 |
| Mouse α-hnRNP A2/B1: Abcam ab5832, dilution 1:1000 |
| Rabbit α-NONO: Bethyl A300-587A-T, dilution 1:1000 |
| Rabbit α-SFPQ: Sigma-Aldrich P2860, dilution 1:1000 |
| Rabbit α-KU80: Cell Signalling #2753, dilution 1:1000 |
| Mouse α-TOPBP1: Santa Cruz Biotechnology sc-271043, dilution 1:1000 |
| Rabbit α-HA: Cell Signalling #2367, dilution 1:1000 |
| Mouse α-Tubulin: Homemade, IFOM, Milan, dilution 1:6000 |
| Mouse α-β-Actin: Abcam ab8226, dilution 1:1000 |
| ***Secondary antibodies (Immunofluorescence)*** |
| AlexaFluor 644 Goat α-Rabbit IgG: ThermoScientific, Life Technologies A11008, dilution |
| AlexaFluor 647 Goat α-Mouse IgG: ThermoScientific, Life Technologies A21235, |
| ***Secondary antibodies (Western Blot)*** |
| Goat α-Rabbit IgG HRP-linked: Cell Signalling #7074, dilution 1:8000 |
| Horse α-Mouse IgG HRP-linked: Cell Signalling #7076, dilution 1:10000 |

SUPPLEMENTARY FIGURES

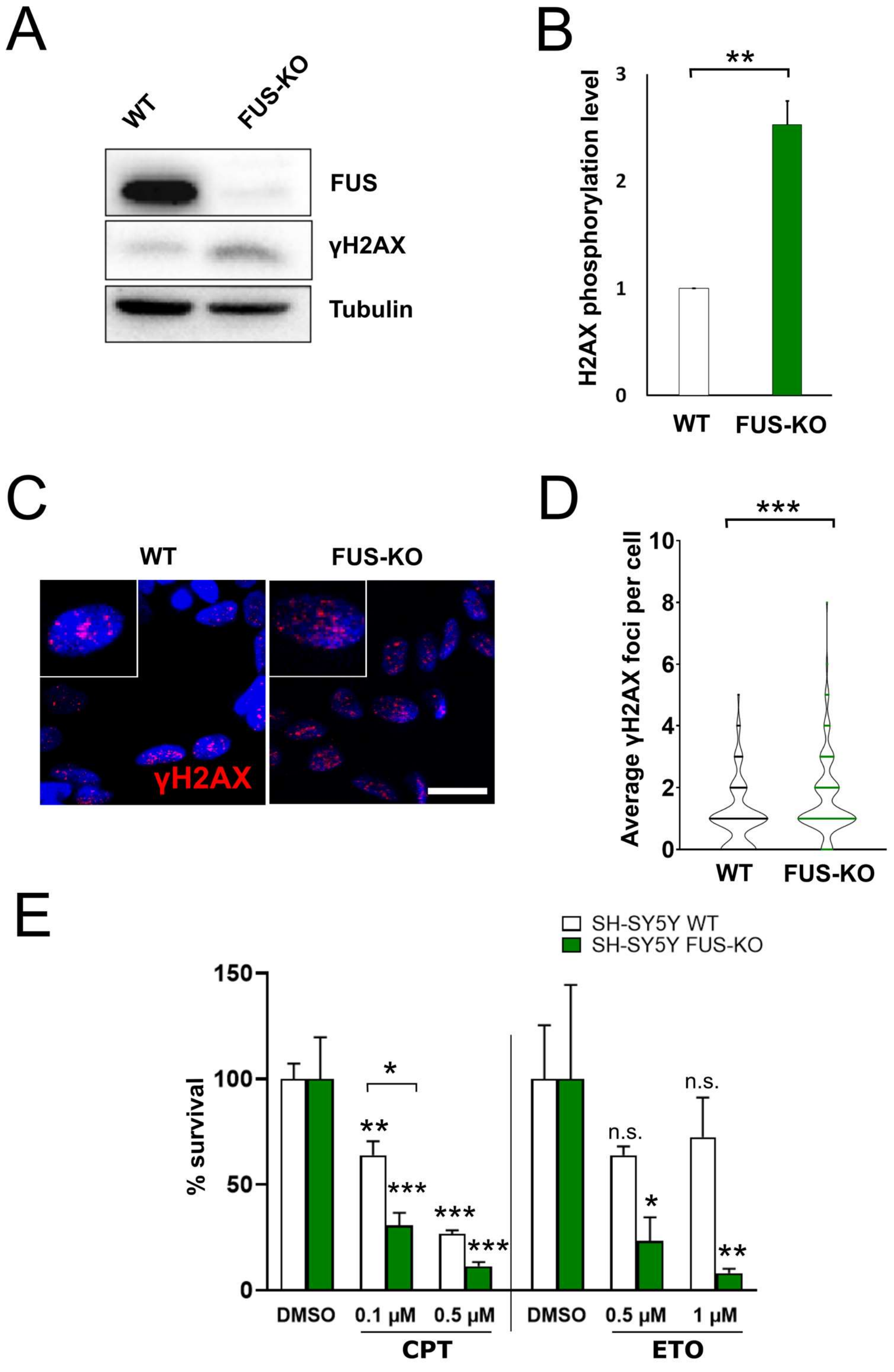

Supplementary Figure 1. **Loss of FUS results in an accumulation of DNA damage and sensitisation to genotoxic insult in SH-SY5Y cells**

A. Total extracts of wild type and FUS-KO SH-SY5Y cells were analysed by Western blotting with anti-FUS and anti-γH2AX antibodies. Tubulin was used as loading control.
B. Quantification of γH2AX protein level of A. Statistical significance was determined using Student's t-test (*** $p < 0.001$).
C. Representative confocal micrographs of γH2AX foci in wild type and FUS-KO SH-SY5Y cells. Cropped single cells are 2x enlarged. Scale bar: 20 μm.
D. Quantification of C. The number of foci per nucleus was counted using ImageJ and plotted as a violin plot. Data from two biological replicates, with 170 cells per replicate. Statistical testing was performed using Student's t-test (*** $p < 0.001$).
E. SH.SY5Y wild type and FUS-KO cell viability assessed by Trypan blue staining upon treatment with increasing concentrations of camptothecin (CPT, 0.1 or 0.5 μM) or etoposide (ETO, 0.5 or 1 μM). Statistical testing was performed using two-way ANOVA followed by Bonferroni post-hoc test (n.s. non-significant, * $p < 0.05$, ** $p < 0.01$ and *** $p < 0.001$).

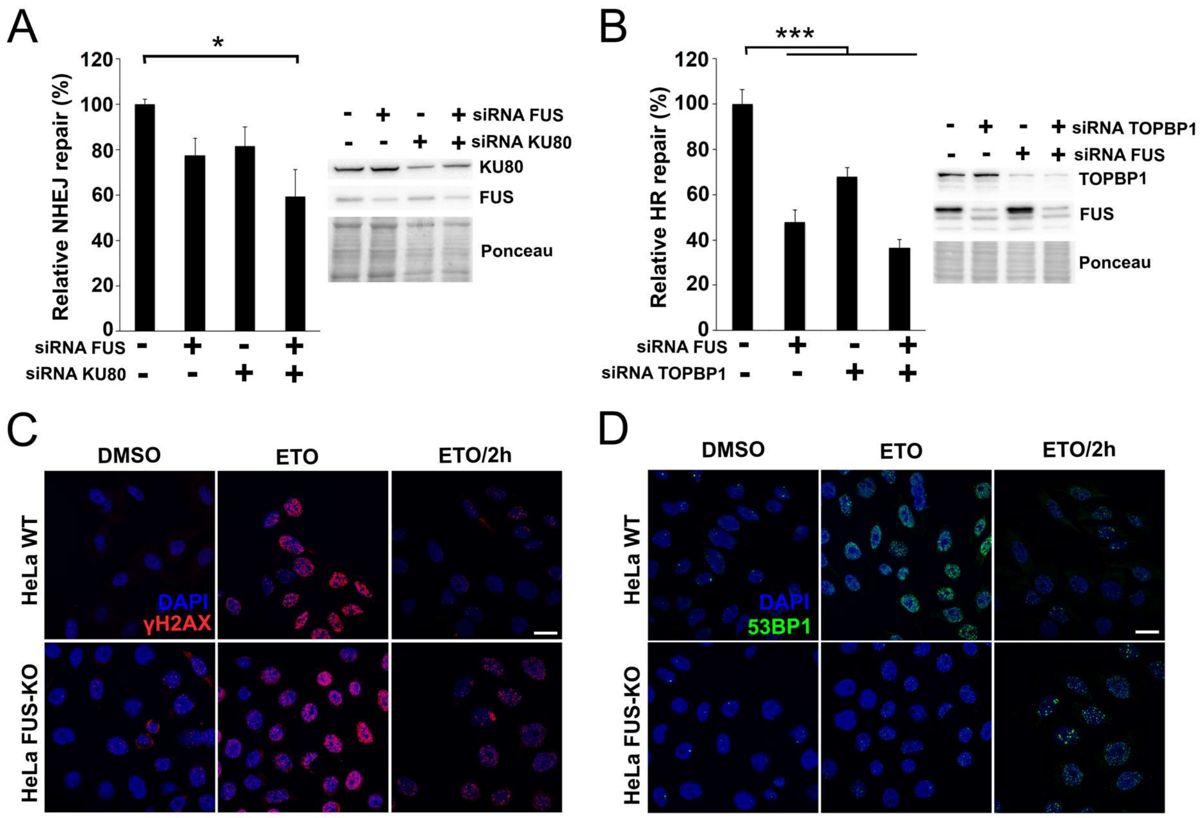

Supplementary Figure 2. **FUS is required for efficient DSB repair and DNA damage foci formation upon genotoxic insult.**

A. DSBs repair efficiency was quantified in U2OS cells containing a stably integrated NHEJ reporter system. Cells were silenced for either FUS, or KU80, or both. Right panel displays Western Blot demonstrating the silencing of the respective proteins. Data are presented as the mean ± SEM (experiments done in triplicate, with at least 10,000 cells analysed per experiment). Statistical significance was assessed by one-way ANOVA, followed by Bonferroni post-hoc test. *$p<0.05$; ***$p<0.001$.

B. DSBs repair efficiency was quantified in U2OS cells containing a stably integrated HR reporter system. Cells were silenced for either FUS, or TOPBP1, or both. Right panel displays Western Blot demonstrating the silencing of the respective proteins. Statistical analysis as in A.

C. HeLa WT and FUS-KO cells were then stained with γH2AX and counterstained with DAPI. Cells were treated with DMSO, etoposide for 1h (ETO) or ETO plus 2 h recovery from ETO treatment (ETO/2h). These are representative figures for graph shown in Figure 2B.

D. HeLa WT and FUS-KO cells were then stained with 53BP1 and counterstained with DAPI. Cells were treated with DMSO, etoposide for 1h (ETO) or ETO plus 2 h recovery from ETO treatment (ETO/2h). These are representative figures for graph shown in Figure 2C.

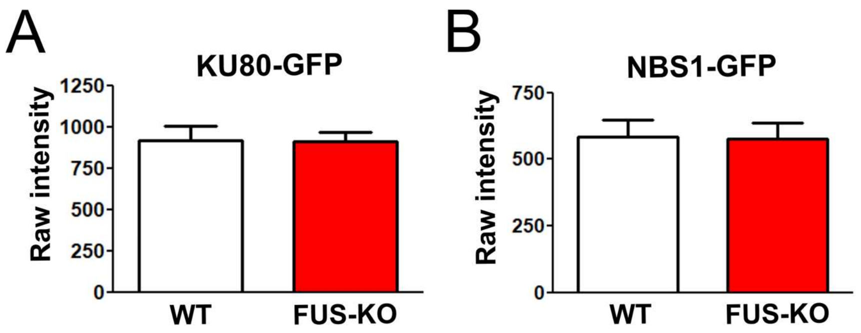

Supplementary Figure 3. **Microirradiation experiments are not influenced by a differential expression of KU80-GFP or NBS1-GFP in WT vs FUS-KO cell lines.**

A. The raw fluorescence intensity of HeLa WT and FUS-KO cells transiently transfected with KU80-GFP was assessed to rule out the possibility that differential expression levels could affect the observed effect in Figure 3A.
B. The raw fluorescence intensity of HeLa WT and FUS-KO cells transiently transfected with NBS1-GFP was assessed to rule out the possibility that differential expression levels could affect the observed effect in Figure 3B.

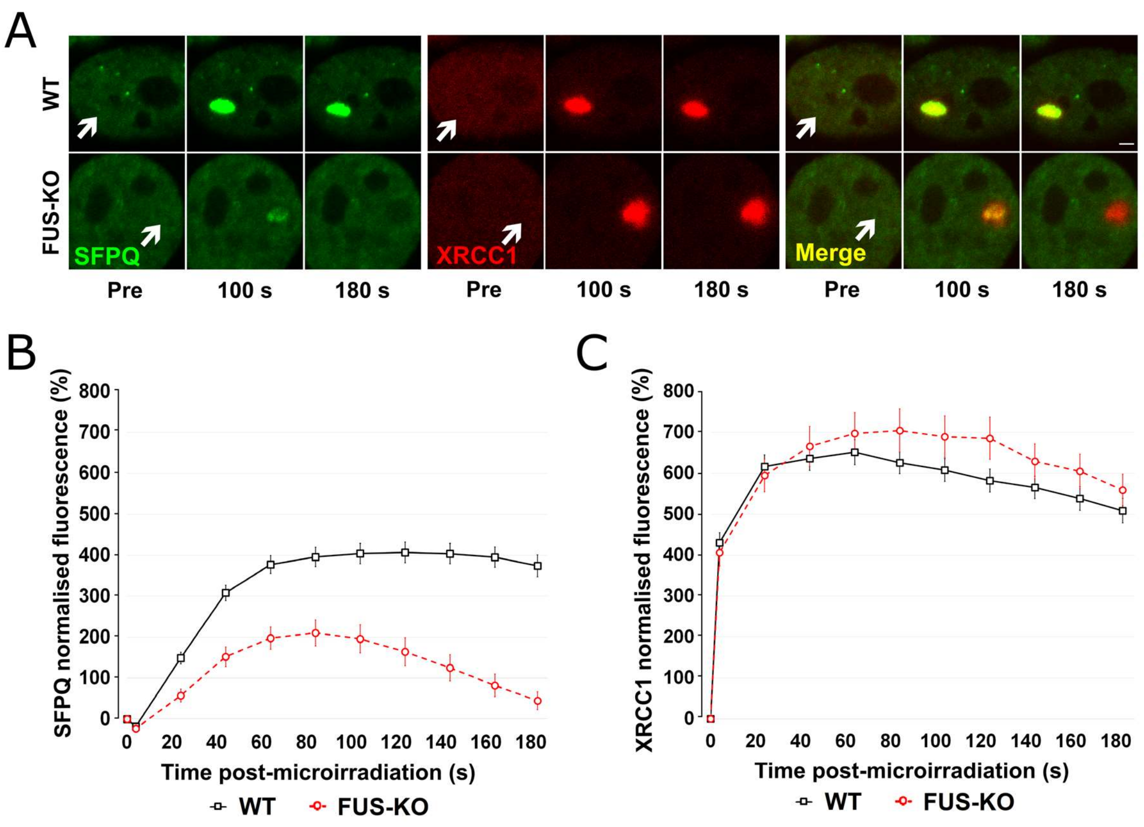

Supplementary Figure 4. **Loss of FUS specifically affects SFPQ but not XRCC1 recruitment to DSB.**

A. WT HeLa cells were transiently co-transfected with SFPQ-GFP and XRCC1-RFP plasmids and submitted to laser microirradiation as described in Materials and Methods. Recruitment of these proteins was assessed for a 3-min period and images were taken every 20 seconds.

B. Recruitment and accumulation of SFPQ, as shown in Figure 4D-E, is severely impaired in FUS-KO cells.

C. Recruitment of XRCC1 is very high (saturated fluorescence signal) and the same for WT and FUS-KO cells.

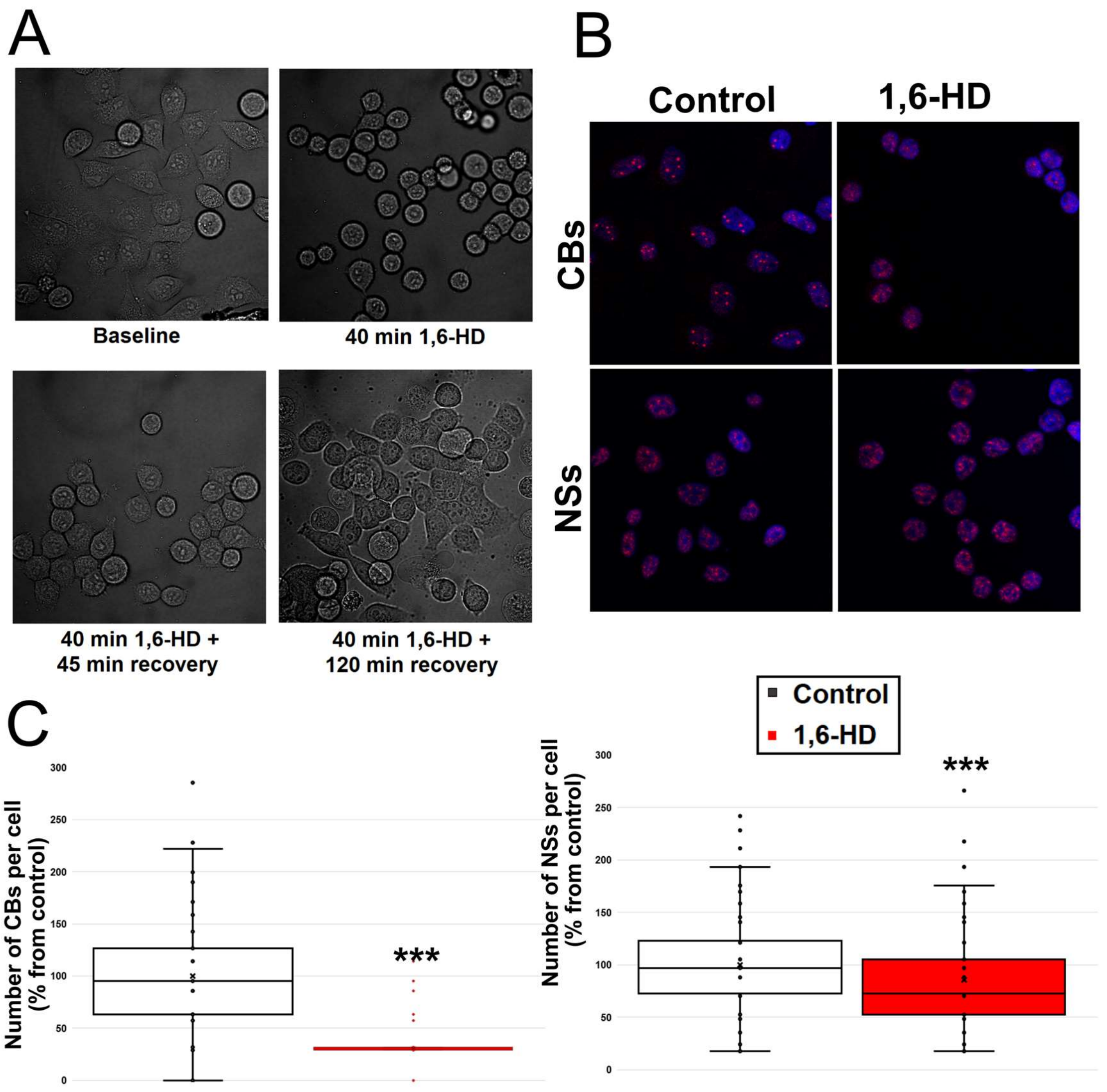

Supplementary Figure 5. **2% 1,6-HD treatment does not irreversibly affect morphology and vitality of cells.**

A. Bright field micrograph of HeLa cells treated with 1.6-HD as described in Material and Methods. Cells were then allowed to recover in alcohol-free medium. Cells returned to a normal morphology within 120 minutes after 1,6-HD withdrawal.

B. Representative images of HeLa WT cells treated or not with 1,6-HD for 30 min and stained for Cajal bodies (CBs, coilin antibody) or Nuclear speckles (NSs, SC-35 antibody).

C. Quantification of Cajal Bodies and Nuclear speckles in untreated and 1,6-HD treated cells. HeLa cells were treated with 2% 1.6-HD for 30 minutes and then stained with DAPI, and either anti-coilin or anti-SC35 antibodies. Quantification was performed as in Material and Methods. Statistical significance was determined using Student's t-test (***$p<0.001$). Experiments were done in duplicate and 200 cells were analysed per experiment.

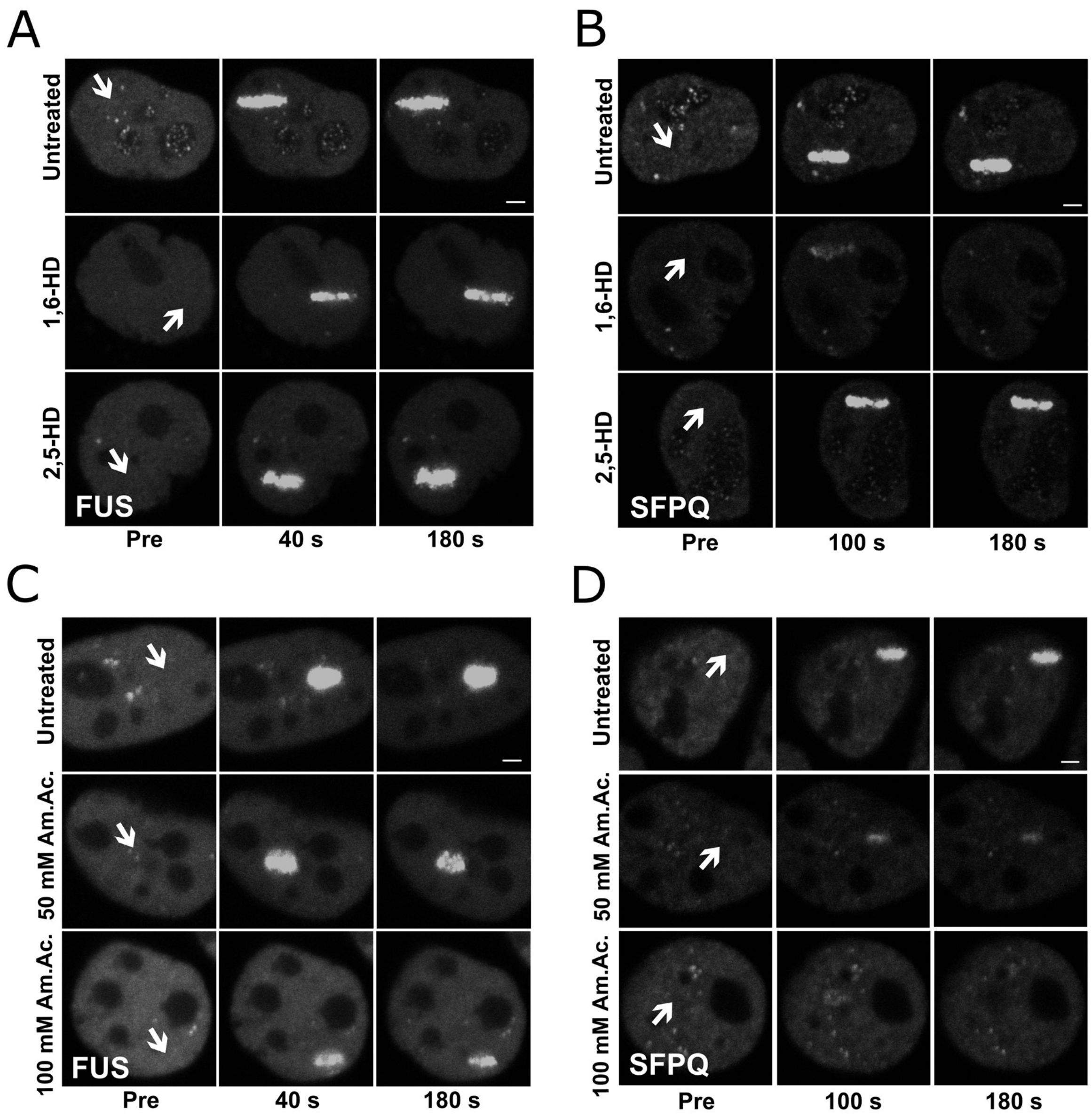

Supplementary Figure 6. **Representative images of the recruitment of SFPQ and FUS in HeLa WT cells treated with 2% 1,6-HD or 2% 2,5-HD.**

A. HeLa WT cells were transfected with FUS-GFP and treated either with 1,6-Hexanediol (1,6-HD) or 2,5-Hexanediol (2,5-HD) prior to laser microirradiation. Scale bar represents 2 µm. Arrows represent irradiated site. These are representative figures for graph shown in Figure 6A.

B. HeLa WT cells were transfected with SFPQ-GFP and treated either with 1,6-Hexanediol (1,6-HD) or 2,5-Hexanediol (2,5-HD) prior to laser microirradiation. Scale bar represents 2 µm. Arrows represent irradiated site. These are representative figures for graph shown in Figure 6B.

C. HeLa WT cells were transfected with FUS-GFP and treated either with 50 mM or 100 mM of Ammonium Acetate (Am. Ac.) prior to laser microirradiation. Scale bar represents 2 μm. Arrows represent irradiated site. These are representative figures for graph shown in Figure 6C.

D. HeLa WT cells were transfected with SFPQ-GFP and treated either with 50 mM or 100 mM of Ammonium Acetate (Am. Ac.) prior to laser microirradiation. Scale bar represents 2 μm. Arrows represent irradiated site. These are representative figures for graph shown in Figure 6D.

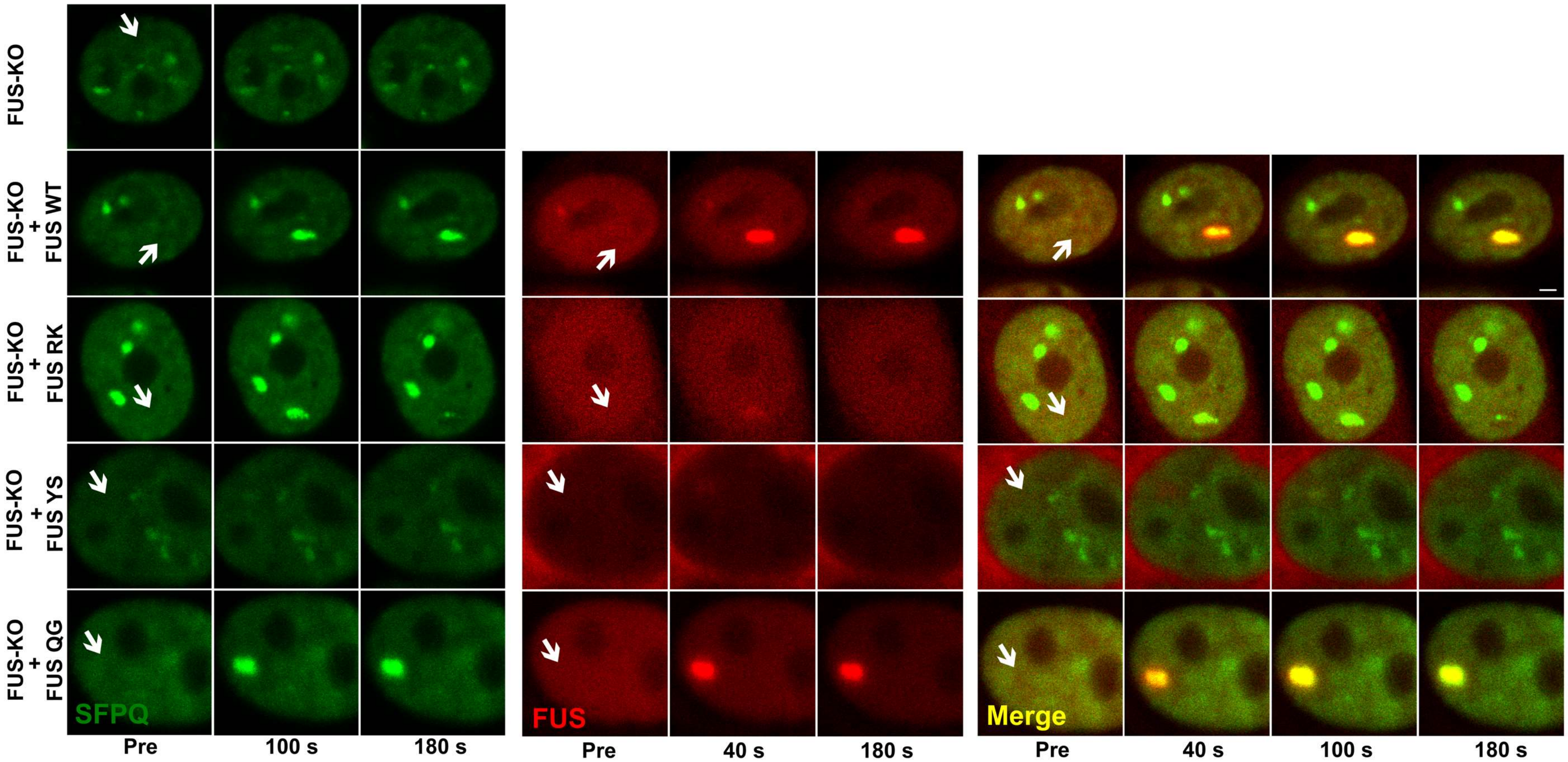

Supplementary Figure 7. **Representative images of the recruitment of SFPQ and FUS in HeLa FUS-KO cells transiently transfected with both SFPQ and a FUS construct (WT or RK, YS or QG mutants).** HeLa FUS-KO cells were transiently transfected with a GFP-SFPQ and co-transfected with one mCherry FUS construct (WT or the mutants RK, YS or QG) and submitted to laser microirradiation. Scale bar represents 2 µm. Arrows represent irradiated site. These are representative figures for graph shown in Figure 5E-F.

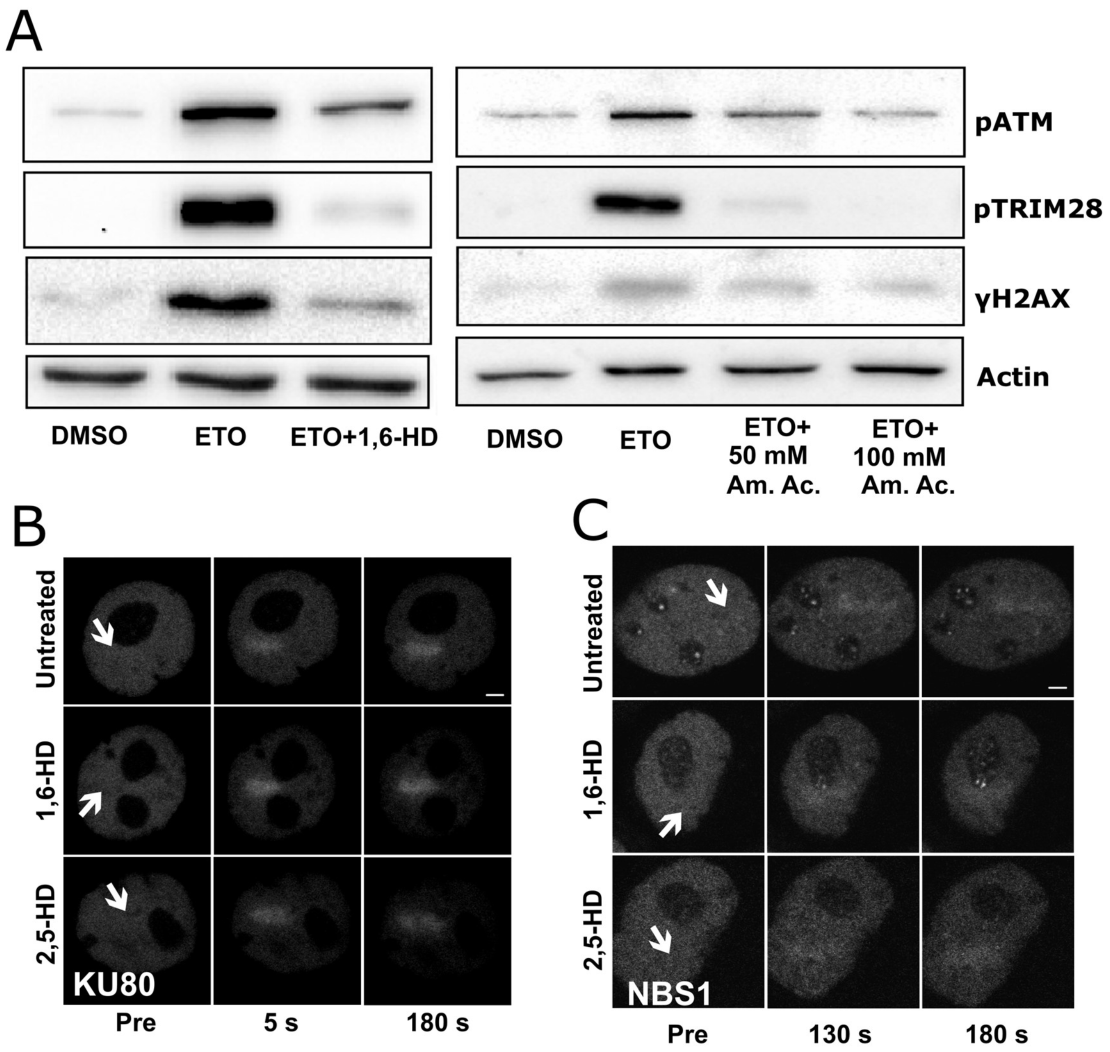

Supplementary Figure 8. **Representative images of the recruitment of KU80 and NBS1 in HeLa WT cells treated with 2% 1,6-HD or 2% 2,5-HD; liquid-liquid phase separation is required for effective DDR signalling.**

A. Western blot analysis of total extracts prepared from HeLa cells that were treated with ETO alone or ETO and 1,6-HD, or ETO and ammonium acetate (Am. Ac., 50 or 100 µM). Phosphorylation of ATM, TRIM28 and H2AX were assessed and actin was used as a normaliser.
B. HeLa WT cells were transfected with KU80-GFP and treated either with 1,6-Hexanediol (1,6-HD) or 2,5-Hexanediol (2,5-HD) prior to laser microirradiation. Scale bar represents 2 µm. Arrows represent irradiated site. These are representative figures for graph shown in Figure 6G.
C. HeLa WT cells were transfected with NBS1-GFP and treated either with 1,6-Hexanediol (1,6-HD) or 2,5-Hexanediol (2,5-HD) prior to laser microirradiation.

Scale bar represents 2 µm. Arrows represent irradiated site. These are representative figures for graph shown in Figure 6H.

**SUPPLEMENTARY VIDEO**

Supplementary Video 1. **SFPQ recruitment in HeLa WT versus FUS-KO cells.** HeLa WT and HeLa FUS-KO cells were transiently transfected with SFPQ-GFP plasmid and laser microirradiated (irradiated area indicated). Video is 10x faster than reality (it shows a window of 180 s). It is possible to observe that the absence of FUS both delays and impairs the recruitment of SFPQ. Scale bar represents 5 µm.
